# *In vivo* inactivation of RAD51-mediated homologous recombination leads to premature aging, but not to tumorigenesis

**DOI:** 10.1101/2022.01.17.476609

**Authors:** Gabriel Matos-Rodrigues, Vilma Barroca, Ali-Akbar Muhammad, Awatef Allouch, Stephane Koundrioukoff, Daniel Lewandowski, Emmanuelle Despras, Josée Guirouilh-Barbat, Lucien Frappart, Patricia Kannouche, Pauline Dupaigne, Eric Le Cam, Jean-Luc Perfettini, Paul-Henri Romeo, Michelle Debatisse, Maria Jasin, Gabriel Livera, Emmanuelle Martini, Bernard S. Lopez

## Abstract

Genetic instability is a hallmark of both cancer and aging. Homologous recombination (HR) is a prominent DNA repair pathway maintaining genomic integrity. Mutations in many HR genes lead to cancer predisposition. Paradoxically, the consequences of mutations in the pivotal HR player, *RAD51*, on cancer development remain puzzling. Moreover, in contrast with other HR genes, RAD51 mouse models are not available to experimentally address the role of RAD51 on aging and carcinogenesis, *in vivo*. Here, we engineered a mouse model with an inducible dominant negative form of RAD51 (*SMRad51*) that suppresses RAD51-mediated HR without stimulating alternative non-conservative repair pathways. We found that, *in vivo* expression of *SMRad51* did not trigger tumorigenesis, but instead induced premature aging. We propose that these *in vivo* phenotypes result from the exhaustion of proliferating progenitors submitted to chronic endogenous replication stress resulting from RAD51-mediated HR suppression. Our data underline the importance of the RAD51 activity for progenitors homeostasis, preventing aging, and more generally for the balance between cancer and aging.

## Introduction

Genetic instability is a hallmark of cancer and aging (1–5). In order to maintain genomic stability, the DNA damage response (DDR) coordinates cell cycle progression and DNA repair. Of note, the activation of the DDR, which is proposed to occur in response to endogenous replication stress, has been observed at the preceding/early stages of cancer and senescence (6–8) suggesting that genetic instability represents an initial step of tumor formation. Moreover, aging and cancer are interrelated and age is considered a risk factor for cancer (2, 3). However, the mechanisms underlying the balance between tissue degeneration *versus* cell transformation that drive aging *versus* cancer upon alteration of DNA repair pathways remain to be fully unraveled (3, 9, 10).

Homologous recombination (HR) is a highly evolutionarily conserved pathway that plays essential roles in genomic plasticity. HR is involved in the repair of DNA double-strand breaks (DSBs) and DNA interstrand crosslinks (ICLs); importantly, it is also involved in the protection of arrested replication forks and the resumption of replication (11, 12). HR is initiated by resection of DNA ends, which generates 3’ single-stranded DNA (ssDNA) tails. These 3’ ssDNA tails are first covered and protected by replication protein A (RPA). Then, BRCA2/PALB2 displace RPA and replace it with RAD51, forming an RAD51-ssDNA filament. This filament promotes homology searching and the invasion of a homologous duplex sequence to form recombination intermediates that will be subsequently resolved. Therefore, the ssDNA-RAD51 filament is the active species of HR and RAD51 thus plays a pivotal role at the critical step of HR.

Because of its importance as replication escort and in genomic stability maintenance, HR is generally considered a tumor suppressor pathway. Consistent with this theory, many HR genes are altered in tumors, both in familial breast and ovarian cancers (13–16) and in sporadic tumors (13, 17). Additionally, many HR genes are mutated in Fanconi anemia (FA) syndrome, a rare autosomal recessive syndrome in which developmental defects are associated with malignancy (13, 18–20). However, in spite of its pivotal role in HR, the implication of RAD51 invalidation in cancer remains debated (13). Several hypothesis have been proposed to account for this paradox (13). One of these hypotheses is based on the fact that mediator/accessory proteins, that are mutated in cancer such as BRCA1/2 or PALB2, promote the loading and the stabilization of RAD51 onto the ssDNA and that the absence of RAD51 on the DNA makes it accessible to alternative non-conservative repair processes such as single strand annealing (SSA) and alternative end-joining A-EJ) that foster genome instability. Therefore, suppression of the mediators/loaders of RAD51, leads not only to the suppression of conservative HR but also to the concomitant stimulation of non-conservative SSA and A-EJ (21–24). This should result in compensation of the decreased viability resulting from HR suppression, but with increased genetic instability. Therefore, this hypothesis raises the question as to whether HR ablation alone would actually be sufficient for oncogenesis, or whether the concomitant stimulation of the non-conservative pathways could be necessary. Addressing this question requires to suppress HR but witghout stimulating SSA and A-EJ.

Mouse models are helpful tools to experimentally address the questions of carcinogenesis and aging, *in vivo*. However, most HR genes are essential, their inactivation leading to embryonic lethality (25–27). Nevertheless, elaborated strategies for partial or tissue-specific HR inactivation have been designed and have confirmed the correlation with cancer development (26). Surprisingly, despite the paramount role of RAD51 in HR, alternative strategies have not been designed to analyze the impact of RAD51 functional disruption *in vivo* (26). Here, we took advantage of an engineered dominant negative form of RAD51 (*SMRad51*), as an experimental tool. One advantage of *SMRad51* is that its expression suppresses RAD51 HR activity without stimulating the alternative mutagenic nonconservative SSA and A-EJ repair pathways (21, 28–32), in contrast with RAD51 or mediators knockdown or with other RAD51 dominant negative forms such as that described in FA (21). Therefore, *SMRad51* is an advantageous implement allowing to precisely focusing on the consequences of HR inhibition itself without interference of the nonconservative SSA and A-EJ repair pathways. We engineered two mouse models with ubiquitous expression of *SMRad51* or, as a control, of exogenous wild-type mouse *Rad51* (exMmRad51) under doxycycline (Dox) induction. This strategy enabled us to overcome the embryonic lethality problem, since HR was inactivated after birth through Dox-mediated induction of *SMRad51*. Thus, our mouse model represents a unique tool to analyze RAD51 functional inactivation *in vivo.* Using this experimental tool, we found that suppression of HR through *SMRad51* expression led to replicative stress, systemic inflammation, progenitor exhaustion, premature aging and reduced lifespan. Remarkably, although *SMRad51* expression induced genetic instability, it did not induce tumor formation.

Although repair deficiency is generally proposed to be a cause of cancer, this work shows that specific RAD51-mediated HR impairment leads primarily to aging rather than oncogenesis. These data shed light on a separation and potential competition rather than the cooperation between these two *in vivo* phenotypes.

## Results

### SMRAD51 suppresses the strand exchange activity of the RAD51-ssDNA filament

*SMRad51* is a yeast/mammalian *Rad51* chimera gene (Fig. 1A) whose ectopic expression suppresses HR in mammalian cells but, importantly, still prevents nonconservative SSA and A-EJ (21, 28–32), unlike RAD51 silencing or suppression of RAD51 loading factors such as for instance BRCA2. To analyze the consequences of specific suppression of the HR function of RAD51 *in vivo*, we developed two mouse models: one bearing Dox-inducible *SMRad51* and one bearing Dox-inducible exogenous wild-type mouse *Rad51* (*exMmRad51*) (Fig.1*B*).

**Fig. 1.**
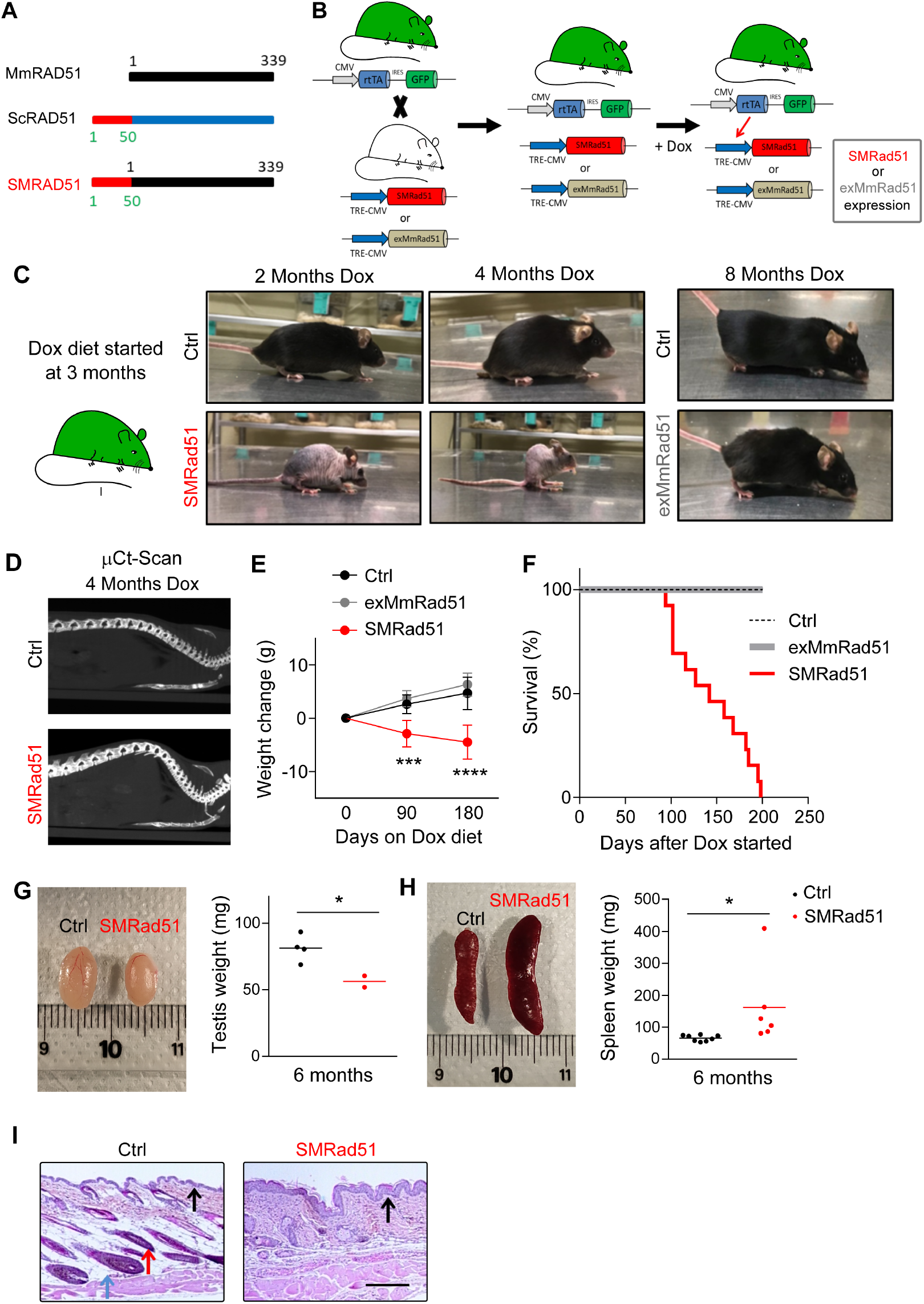
Functional inactivation of RAD51 leads to premature aging. **A.** Schematic representation of SMRAD51. SMRAD51 is a chimeric construct composed of the full-length wild-type mouse MmRAD51 (black box) fused to the non-evolutionarily conserved N-terminal 55 amino acids (red box) of *Saccharomyces cerevisiae* Rad51 (ScRad51). **B.** Schematic representation of SMRad51 and exMmRad51 mouse model generation. Two mouse models, one with SMRad51 and the other with exMmRad51 (HA-tagged) under the control of the Dox-inducible promoter (TRE-CMV), were designed. The induction of transgene expression by Dox requires the rtTA transcription activator. The two transgenic mouse models were thus crossed with a mouse bearing rtTA (and GFP) at the ubiquitous expression locus *Rosa26*. Expression of the transgene (*SMRad51* or *exMmRad51*) resulted from exposure of the *rtTA+;SMRad51+* or *rtTA+;exMmRad51+* models to Dox. The *rtTA-;SMRad51+* and *rtTA-;exMmRad51+* mice did not express the transgenes, even in the presence of Dox, and were thus convenient controls. **C.** Experimental design and representative photos of Ctrl, exMmRad51 and SMRad51 mice fed a Dox-containing diet for 2, 4 or 8 months. **D.**μCt scan images illustrating the spines of Ctrl and SMRad51 adult mice fed a Dox-containing diet for 4 months. **E.** Quantitative analysis of body weight changes from the beginning of Dox treatment in Ctrl mice (n=9, 3 months; n=7, 6 months), exMmRad51 mice (n=5, 3 months; n=5, 6 months) and SMRad51 mice (n=10, 3 months; n=5, 6 months) fed a Dox-containing diet for 3 and 6 months. **F.** Survival curves of Ctrl, exMmRad51 and SMRad51 mice fed a Dox-containing diet. Illustrative images (left) and quantification (right) of testis weights in Ctrl and SMRad51 adult mice fed a Dox-containing diet for 6 months. **G.** Illustrative images (left) and quantification (right) of testis weights in Ctrl and SMRad51 adult mice fed a Dox-containing diet for 6 months. **H.** Illustrative images (left) and quantification (right) of spleen weights in Ctrl and SMRad51 adult mice fed a Dox-containing diet for 6 months. **I.** Representative pictures of skin histology for Ctrl and SMRad51 mice fed a Dox-containing diet for 6 months. Red arrow: capillary bulb, the density of which was decreased in SMRad51-expressing mice. Blue arrow: hypodermis in a Ctrl mouse, the thickness of which was decreased in *SMRad51*-expressing mice. Black arrow: epidermis, the thickness of which was increased in SMRad51 mutants. Statistical analysis: **(E)** Two-way ANOVA followed by Sidak’s post-test. *** p<0.001; ****p<0.0001. **(G-I)** Student’s t-test. *p<0.05.

In mouse embryonic fibroblasts (MEFs) derived from our mouse models, Dox-induced *SMRad51* expression decreased homology-directed gene targeting efficacy, increased sensitivity to DNA-damaging agents, and increased genomic instability, compared to the absence of Dox (*SI Appendix*, Fig. S1*A-D*). All these phenotypes indeed reflect HR deficiency.

To further elucidate the molecular basis of the SMRAD51 dominant negative effect, we purified SMRAD51 and mouse wild-type MmRAD51 proteins (*SI Appendix*, Fig. S2*A*). The D-loop *in vitro* assay revealed that adding SMRAD51 to MmRAD51 inhibited its strand invasion activity (*SI Appendix*, Fig. S2*B* and S2*C*), accounting thus for the dominant negative effect observed in living cells. Transmission electron microscopy (TEM) revealed that the SMRAD51 protein was able to coat DNA *in vitro* (*SI Appendix*, Fig. S2*D*). Note that 100% of the DNA molecules were coated by either MmRAD51 or SMRAD51. These data indicate that SMRAD51 inhibits HR by poisoning the RAD51-ssDNA filament strand exchange activity, thus inhibiting the central HR activity.

Importantly, SMRAD51 formed radiation-induced nuclear foci with assembly/disassembly kinetics similar to those of exMmRAD51 as well as endogenous MmRAD51 (*SI Appendix*, Fig. S2*E* and S2*F*), confirming the in vitro TEM data showing that SMRAD51 can bind DNA, and that it is processed similarly to wild-type endogenous MmRAD51, in living cells.

Collectively, these data show that SMRAD51 inhibits the strand invasion activity of the RAD51/ssDNA filament, which accounts for its dominant negative effect on HR, and that the consequences of SMRAD51 expression result from RAD51-ssDNA filament activity disruption rather than unscheduled prolonged DNA occupancy by the SMRAD51 protein. Note that the ssDNA occupancy by SMRAD51 also account for its capacity to prevent accessibility to non-conservative repair, as proposed (21).

### Suppression of RAD51 activity leads to aging rather than carcinogenesis

To analyze the *in vivo* consequences of specific inhibition of RAD51-HR activity, we induced the expression of *SMRad51* or *exMmRad51* in *rtTA+;SMRad51+* and *rtTA+;exMmRad51+* mice, respectively. These mice were compared to Dox-exposed control littermates (*rtTA-;SMRad51+* and *rtTA-;exMmRad51+*) that bore the transgenes but not the transcription activator *rtTA* and thus did not express the transgenes in the presence of Dox.

We fed 3-month-old adult mice *ad libitum* with Dox-containing food (Fig. 1*C*). After approximately 6 weeks of Dox exposure, SMRad51, but not exMmRad51, mice started to show features of premature aging, including reduced activity, hair loss, intermittent priapism and abnormal posture with protuberance of the upper back (Fig. 1*C*; *SI Appendix*, Table S1). X-ray imaging by micro-computed tomography (μCT) after 4 months of Dox treatment revealed that these back changes were associated with curvature of the spine, consistent with kyphosis (Fig. 1*D*, Video S1 and S2). Prolonged expression of *SMRad51*, but not *exMmRad51*, decreased both body weight and life span (Fig. 1*E* and 1*F*). Collectively, these phenotypes support the induction of premature aging in *SMRad51*-expressing mice.

To evaluate tissue morphological modifications caused by *SMRad51* expression, we performed anatomical and histopathological analyses. After 6 months of Dox treatment, we observed testis size reductions (Fig. 1*G*) and splenomegaly, as shown by increased spleen sizes (Fig.1*H*). Splenomegaly is a feature of systemic inflammation and aging in mice (33). We then performed histopathological analyses of different tissues from mice fed a Dox-supplemented diet for 6 months (*SI Appendix*, Fig. S3*A*). These analyses revealed edematous alveolitis in the lungs of *SMRad51*-expressing mice (*SI Appendix*, Fig. S3*B*), consistent with induction of inflammation. Moreover, *SMRad51* expression decreased capillary bulb density and led to hyperplasia in the epidermis (Fig. 1*I*). Subcutaneous skin fat layer thickness was reduced in *SMRad51*-expressing mice compared to control mice (Fig. 1*I*). Coherently with an aging phenotype, reduction in subcutaneous fat has been observed in aged and prematurely aged mice (34). Similar phenotypes are observed in mice with skin-specific CRE-LOX-mediated inactivation of *Brca1* (35). Given that BRCA1 and RAD51 share roles in HR, this finding suggests that the phenotypes observed here actually resulted from HR inactivation *in vivo*. The other tissues that we analyzed did not present major histological modifications (*SI Appendix*, Fig. S3*A*). Remarkably, tumors were detected in none of the animals and none of the different tissues by the wide anatomical and histopathological analysis; only one mouse had a precancerous lesion in the skin (*SI Appendix*, Fig. S3*A*). Altogether, these data show that *SMRad51* expression leads to premature aging but not to increased tumorigenesis.

Since we observed morphological modifications associated with inflammation in several organs, we evaluated whether *SMRad51* expression leads to a systemic inflammatory response. We measured the levels of cytokines in the serum of SMRad51 and control mice after three months of Dox exposure. Using cytokine arrays, we showed that among the 111 proteins analyzed, 29 were upregulated in SMRad51 mice, most of which were proinflammatory factors (Fig. 2). These data show that functional disruption of RAD51 leads to a systemic inflammatory response in adult mice.

**Fig. 2.**
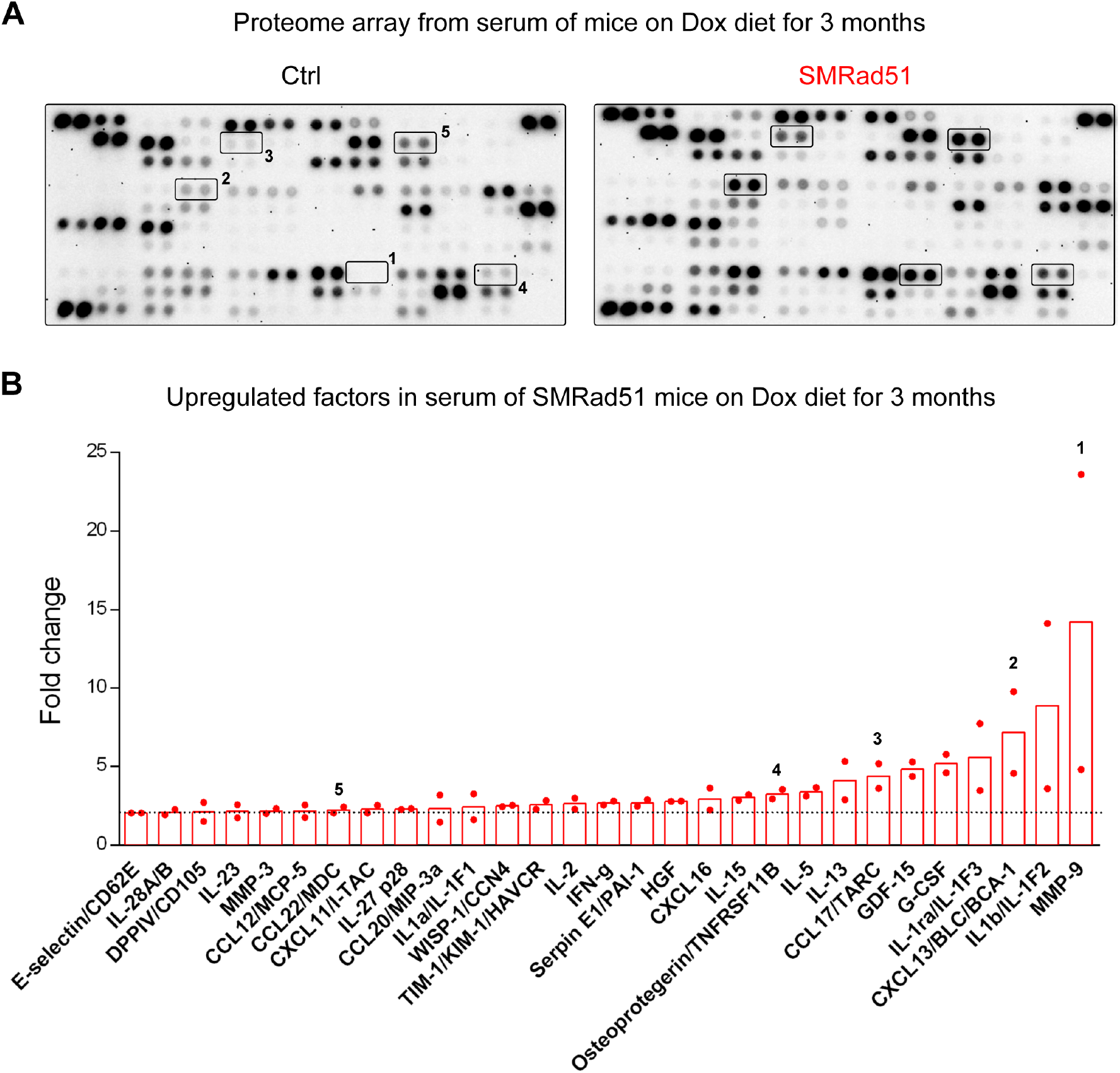
Expression of *SMRad51* in adult mice increases serum levels of proinflammatory cytokines. **(A)** Representative images of proteome array membranes of Ctrl and SMRad51 mice fed a Dox-containing diet for 3 months. **(B)** Graphical representation of the fold changes in upregulated factors in SMRad51 mouse serum compared to Ctrl mouse serum. Each dot in the graph represents a different biological replicate, and the horizontal dashed line marks a 2-fold change. *SI Appendix*, Table S3 shows the quantification for the two independent biological samples of each group. In **(A)** and **(B)**, the numbers illustrate 5 different duplicate protein dots in the membranes and their quantification in two different experiments.

### Suppression of RAD51-mediated HR alters progenitor homeostasis

We observed the presence of hematopoietic cells in the spleens of SMRad51 mice after 6 months of Dox exposure, revealing extramedullary hematopoiesis (*SI Appendix*, Fig. S4). Splenomegaly associated with extramedullary hematopoiesis is a common feature of aged and prematurely aged mice and is a compensatory mechanism triggered by bone marrow progenitor exhaustion (33, 36, 37).

Then, we investigated whether these changes in spleen histology were associated with alterations in hematopoiesis. First, we analyzed the blood composition of mice with and without *SMRad51* expression. The proportions of red blood cells (RBCs) and platelets (PLTs), but not those of white blood cells (WBCs), in the blood were decreased in SMRad51 mice compared to controls after three months of Dox treatment (Fig. 3*A*). Next, we evaluated whether these changes correlated with hematopoietic changes in the bone marrow. Although *SMRad51* expression did not reduce the global content of bone marrow stem cells (Lineage [Lin]^−^Sca-1^+^-c-Kit^+^-Fkl2^−^ or LSK, Flk2^−^), it reduced the proportion of common lymphocyte progenitors (CLPs; Lin^−^ Sca-1^−^c-Kit^+^IL7R^+^) and B-cells (B220^+^). These data indicate that SMRAD51 alters hematopoiesis by diminishing the expansion of progenitor cells (Fig. 3*B*). Altogether, our data show that expression of *SMRad51* disrupts blood cell production, leading to thrombocytopenia (a reduction in PLTs) and anemia (a reduction in RBCs) associated with compromised hematopoiesis.

**Fig. 3.**
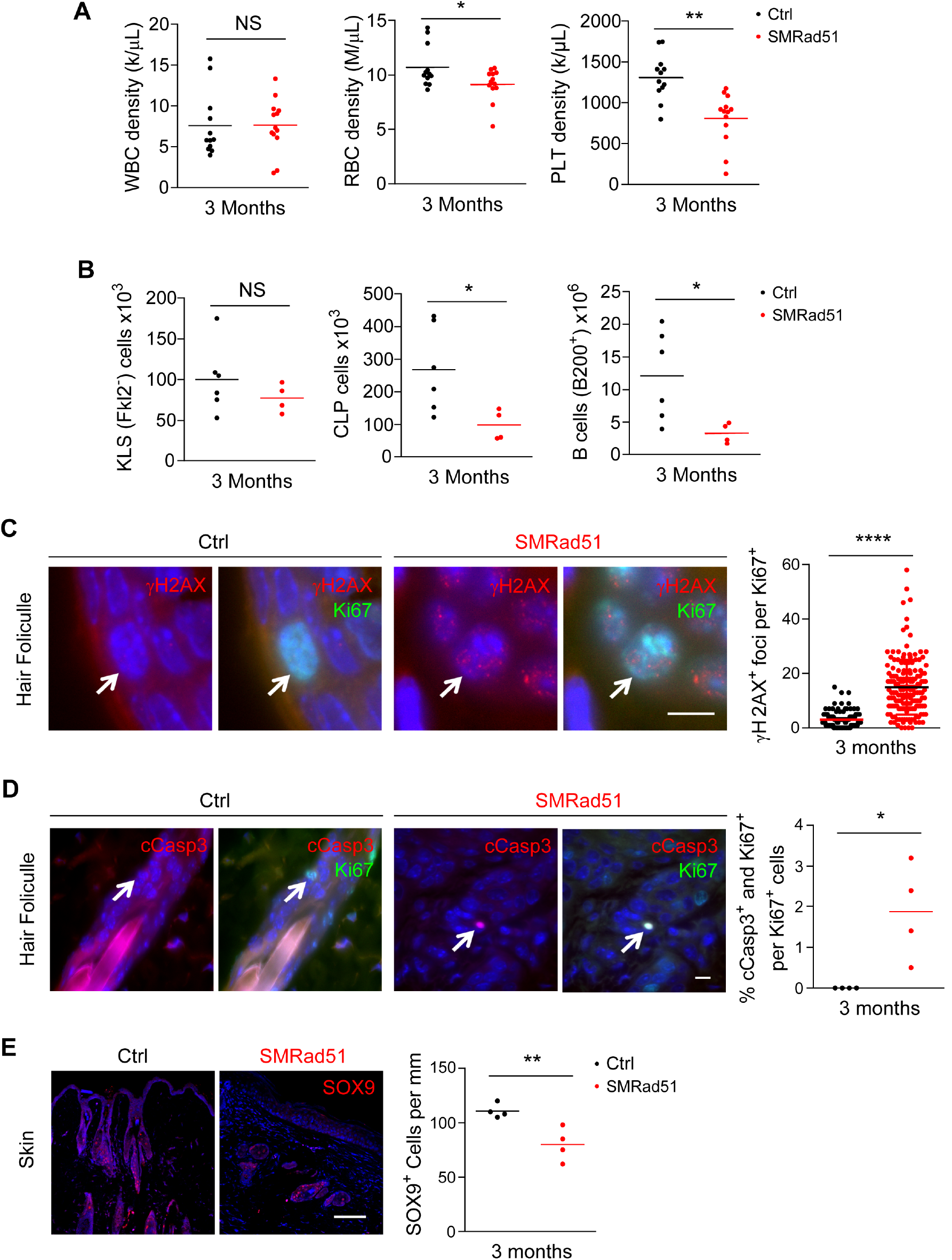
SMRad51 expression disrupts progenitor homeostasis. **A.** Analysis of the density of WBCs, RBCs and PLTs in the blood of Ctrl and SMRad51 adult mice 3 months after Dox treatment started. **B.** Analysis of the proportion of hematopoietic stem cells (KLS cells, FLK2-), CLPs and B-lymphocytes (B-cells, B220+) in the bone marrow of Ctrl and SMRad51 adult mice 3 months after Dox treatment started. **C.** Representative images (left) and quantification (right) of γ H2AX foci in Ki67^+^ (white arrow) capillary bulb cells in skin sections of Ctrl and SMRad51 mice fed a Dox-containing diet for 3 months. **D.** Representative images (left) and quantification (right) of cCas3^+^ and Ki67^+^ (white arrow) capillary bulb cells in skin sections of Ctrl and SMRad51 mice fed a Dox-containing diet for 3 months. **E.** Representative images (left) and quantification (right) of SOX9^+^ cells in skin sections of Ctrl and SMRad51 mice fed a Dox-containing diet for 3 months. Statistical analysis: **(c)** Mann-Whitney test and **(A, B, C, D, E)** Student’s t-test. *p<0.05; **p<0.01; ****p<0.0001. Scale bars: **(C, D)** 10 μm and **(e)** 100 μm.

Progenitor exhaustion is a prime cause of aging (3). Since SMRAD51 suppresses HR, which is a prime DNA damage repair pathway, we evaluated whether DNA damage and apoptosis were correlated with progenitor loss upon *SMRad51* expression. Double immunostaining for Ki67 (a progenitor marker) and γH2AX (a DNA damage marker) showed an increase in DNA damage in capillary bulb progenitors in the skin of *SMRad51*-expressing mice after 3 months of Dox treatment (Fig. 3*C*). In addition, cleaved caspase-3 (cCasp3) staining revealed an increase in apoptosis of capillary bulb progenitors (Fig. 3*D*). The increases in DNA damage and apoptosis were also associated with reductions in the numbers of SOX9^+^ cells, the progenitors of hair follicles, in mice expressing *SMRad51* for 3 months (Fig. 3*E*). Taken together, these data support the idea that apoptosis triggered by chronic DNA damage is an underlying cause of progenitor loss in mice expressing *SMRad51*, thus accounting for the premature aging phenotypes.

### Disruption of RAD51 HR activity strongly compromises growing mice viability

While suppression of RAD51 activity did not lead to tumorigenesis in adult mice, we considered it possible that HR inactivation in young growing mice, which bear more dividing cells, could be necessary for tumor development in adults. We therefore expressed SMRAD51 in young growing mice and observed that it induced phenotypes much more rapidly than in adult mice without induction of tumorigenesis. Expression of SMRAD51 resulted in growth arrest (scored on the basis of body weight), hair loss and death as soon as day 7 of Dox exposure in mice at postnatal days 12-14 (P12-P14) (Fig. 4*A-D).* In contrast, *exMmRad51* expression did not cause hair loss or affect body weight or mouse survival, even after prolonged Dox treatment for up to 3 weeks (Fig. 4*A-D*).

**Fig. 4.**
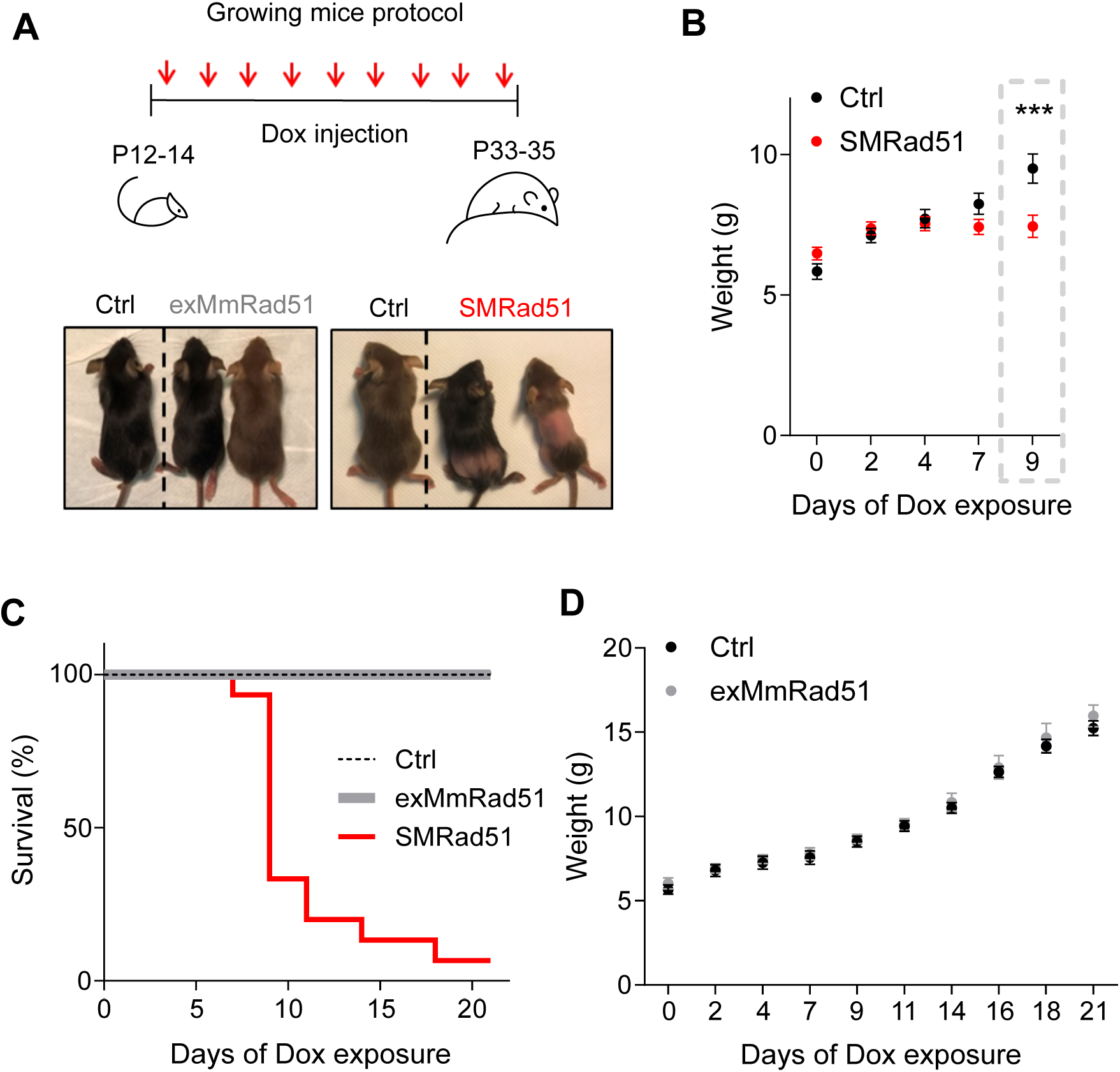
*SMRad51*, but not *exMmRad51*, expression leads to rapid death in young growing mice. **A.** Representative scheme of the experimental designs for exMmRad51 and SMRad51 expression in growing mice (top) and representative photos (bottom) of Ctrl, exMmRad51 and SMRad51 growing mice after 12 days of Dox treatment. **B.** Body weight measurements in SMRad51 (n=21) and Ctrl (n=15) littermate mice treated with Dox. **C.** Survival curve from P12/14 to P33/35 in Ctrl (n=32), exMmRad51 (n=6) and SMRad51 (n=15) mice treated with Dox. **D.** Body weight measurements in exMmRad51 (n=6) and Ctrl (n=10) littermate mice treated with Dox. Statistical analysis: **(B)** Two-way ANOVA followed by Sidak posttest. *** p<0.001. The error bars represent the ±SEM. **(C)** Student’s t-test. * p<0.05.

The presence of the SMRAD51 protein was observed in several tissues (*SI Appendix*, Fig. S5*A*). Histopathological analysis of young mice that survived 12 days of *SMRad51* expression (from P12-14 to P24-26) revealed morphological alterations in highly proliferative tissues, such as the skin and testes (Fig. 5*A*, 5*B* and *SI Appendix*, Fig. S5*B*). In accordance with our data for adult mice, *SMRad51* expression in the skin led to capillary bulb atrophy, subcutaneous fat loss, and epidermal hyperplasia (Fig. 5*B*). Moreover, the bone marrow of *SMRad51*-expressing mice exhibited a decreased cell number (*SI Appendix*, Fig. S5*C*). More specifically, *SMRad51* expression decreased progenitor cell populations in highly proliferative tissues: SOX9^+^ cells in the skin (Fig. 5*B*), PLZF^+^ cells in the testis (Fig. 5*D*), and Lin^−^Sca-1^+^c-Kit^+^ (LSK) stem cells/progenitors in the bone marrow (Fig. 5*E*). Thus, in growing mice, SMRAD51 affects progenitor homeostasis in different highly proliferative tissues.

**Fig. 5.**
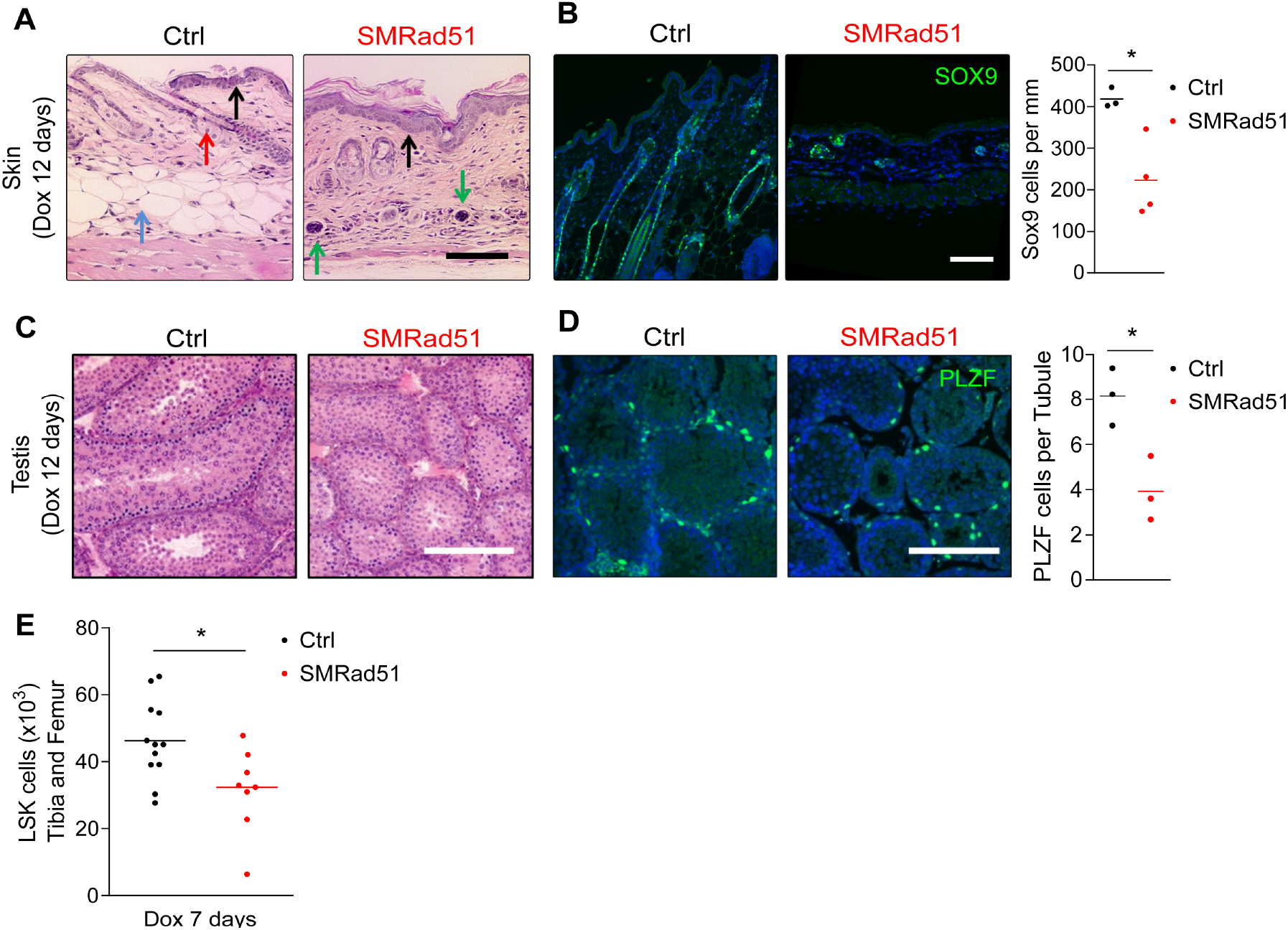
*SMRad51* expression in young growing mice decreases progenitor numbers in highly proliferative tissues. **A.** Representative pictures of histological analysis of skin from Ctrl and SMRad51 mice treated with Dox. Red arrow: capillary bulb, the density of which was decreased in SMRad51-expressing mice. Blue arrow: hypodermis, the thickness of which was decreased in SMRad51-expressing mice. Black arrow: epidermis, the thickness of which was increased in SMRad51 mutants. Green arrow: inflammatory infiltrates. **B.** Representative pictures and quantification of capillary bulb progenitors stained for SOX9 in skin sections of Ctrl and SMRad51 mice treated with Dox. **C.** Representative pictures of Ctrl and SMRad51 testis sections stained with H&E. **D.** Representative images (left) and quantification (right) of testis sections immunostained for PLZF in Ctrl and SMRad51 mice after 12 days of Dox treatment. **E.** Numbers of bone marrow stem cells (LSK, Lin-Sca1+c-kit+) in the tibias and femurs of Ctrl and SMRad51 mice treated for 7 days with Dox. Statistical analysis: **(B, D and E)** Student’s t-test. * p<0.05. Each point represents a biological replicate. Scale bar: 100 μm.

### Disruption of RAD51 HR activity induces inflammation in growing mice

Histopathological analysis of the skin revealed the presence of inflammatory infiltrates after 12 days of *SMRad51* expression in growing mice (Fig. 5*A*). Immunolocalization analysis revealed an increase in T-lymphocyte infiltration (as indicated by CD3 staining; Fig. 6*A*) and monocyte/macrophage aggregates (as indicated by F4/80 staining) in skin samples (Fig. 6*B*). Real-time RT-PCR showed that SMRAD51 stimulated the expression of proinflammatory cytokines in the skin (Fig. 6*C*). Thus, *SMRad51* expression leads to a tissue inflammatory response *in vivo*. These data are consistent with the induction of systemic inflammation observed in adult mice (see Fig. 2). We next investigated whether *in vivo* expression of *SMRad51* also induces a systemic inflammatory response in young growing mice. No differences in the density of WBCs, RBCs or hemoglobin (HGB) were observed between SMRad51-expressing and control growing mice (7 days of Dox treatment) (*SI Appendix*, Fig. S6). However, *SMRad51* expression decreased B-lymphocyte numbers and increased monocyte numbers (Fig. 6*D*), in agreement with the induction of inflammation (38, 39). To further analyze the systemic effects of *SMRad51* expression, we performed a serum cytokine array analysis (Fig. 6*E*). Proinflammatory factors that were upregulated in the serum of SMRad51 adult mice, namely, Lipocalin-2/NGAL, CCL17/TARC, E-selectin/CD62A and CCL22/MDC, were also among the most upregulated factors in the serum of young mice (compare Fig. 6*E* and Fig. 2). These data show that *SMRad51* expression leads to a systemic proinflammatory response in both adult and growing mice.

**Fig. 6.**
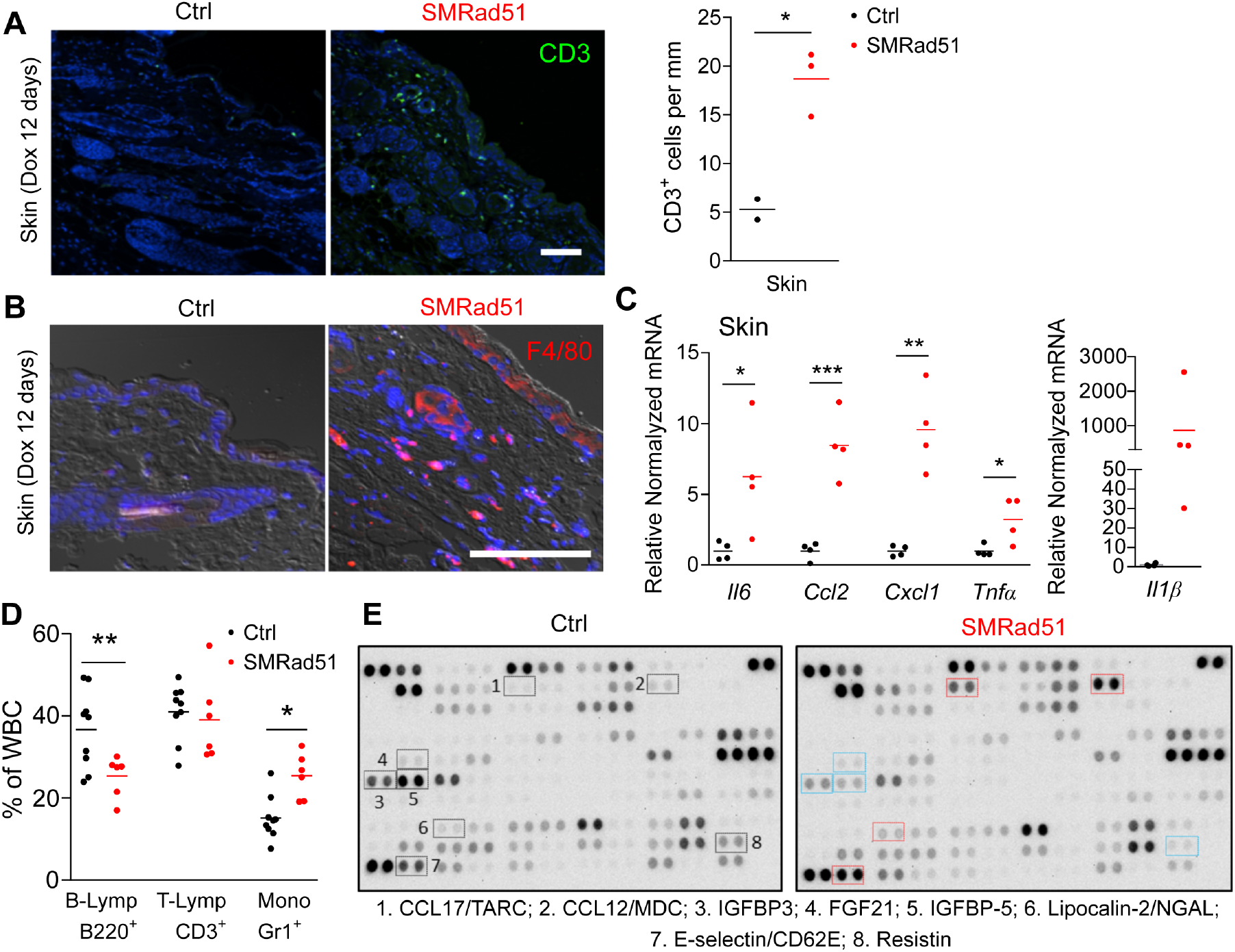
Functional inhibition of *SMRad51* expression induces inflammation in young growing mice. **A.** Representative images (left panels) and quantification (right panel) of CD3^+^ cells (T-lymphocytes) in skin sections of Ctrl and SMRad51 mice treated with Dox for 12 days. **B.** Representative images of staining for F4/80 (a monocyte/macrophage marker) in skin sections of Ctrl and SMRad51 mice treated with Dox for 12 days. **C.** Real-time RT-PCR analysis of Ctrl and SMRad51 skin samples. **D.** WBC proportions in growing mice after 7 days of Dox treatment. B-lymphocytes (B-Lymp) were revealed by B220 staining. T-lymphocytes (T-Lymp) were revealed by CD3 staining. Monocytes (Mono) were revealed by Gr1 staining. **E.** Representative image of cytokine array analysis using serum from SMRad51 and Ctrl growing mice after 7 days of Dox treatment. The selected proteins highlighted in blue were downregulated (SMRad51 vs Ctrl), and those highlighted in red were upregulated (SMRad51 vs Ctrl). *SI Appendix*, Table S4 shows the quantification results for two independent biological samples of each group. Statistical analysis: **(A, C, D)** Student’s t-test. *p<0.05; **p<0.01; ***p<0.001. Each point represents a biological replicate. Scale bar: 100 μm.

### SMRAD51 affects replication dynamics

Our data suggest that replication dynamics might be corrupted in *SMRad51-*expressing cells given (i) the pronounced effect of *in vivo SMRad51* expression on proliferative tissues; (ii) the fact that growing mice, in which many tissues are proliferating, are more sensitive than adult mice; and (iii) the role of HR in arrested replication fork resumption. In addition, the connections between replication stress and inflammation that are now clearly established could account for the observed inflammation (40–43). Therefore, we evaluated the impact of SMRAD51 on replication dynamics in our mouse models.

In the primary MEFs, the expression of *SMRad51*, but not *exMmRad51*, increased the number of pCHK1 (S317) and γH2AX foci, indicating spontaneous activation of the DDR in response to endogenous replication stress (Fig. 7*A* and 7*B*). Molecular combing revealed that SMRAD51, but not exMmRAD51, decreased the velocity of DNA replication and concomitantly increased the frequency of asymmetric labeling, indicating the accumulation of arrested replication forks (Fig. 7*C-E*). This finding suggested that resumption of replication at the arrested forks, which normally involves RAD51-mediated strand exchange activity for template switching, was defective. To analyze the impact of *SMRad51* expression on fork restart, we performed a DNA spreading analysis (Fig. 7*F*) after blocking replication with hydroxyurea (HU), a ribonucleotide reductase inhibitor that generates replication stress through nucleotide pool exhaustion. SMRAD51 decreased replication restart efficiency after HU release (Fig. 7*F*), consistent with the inhibition of strand exchange by SMRAD51 (see *SI Appendix*, Fig. S2*A*).

**Fig. 7.**
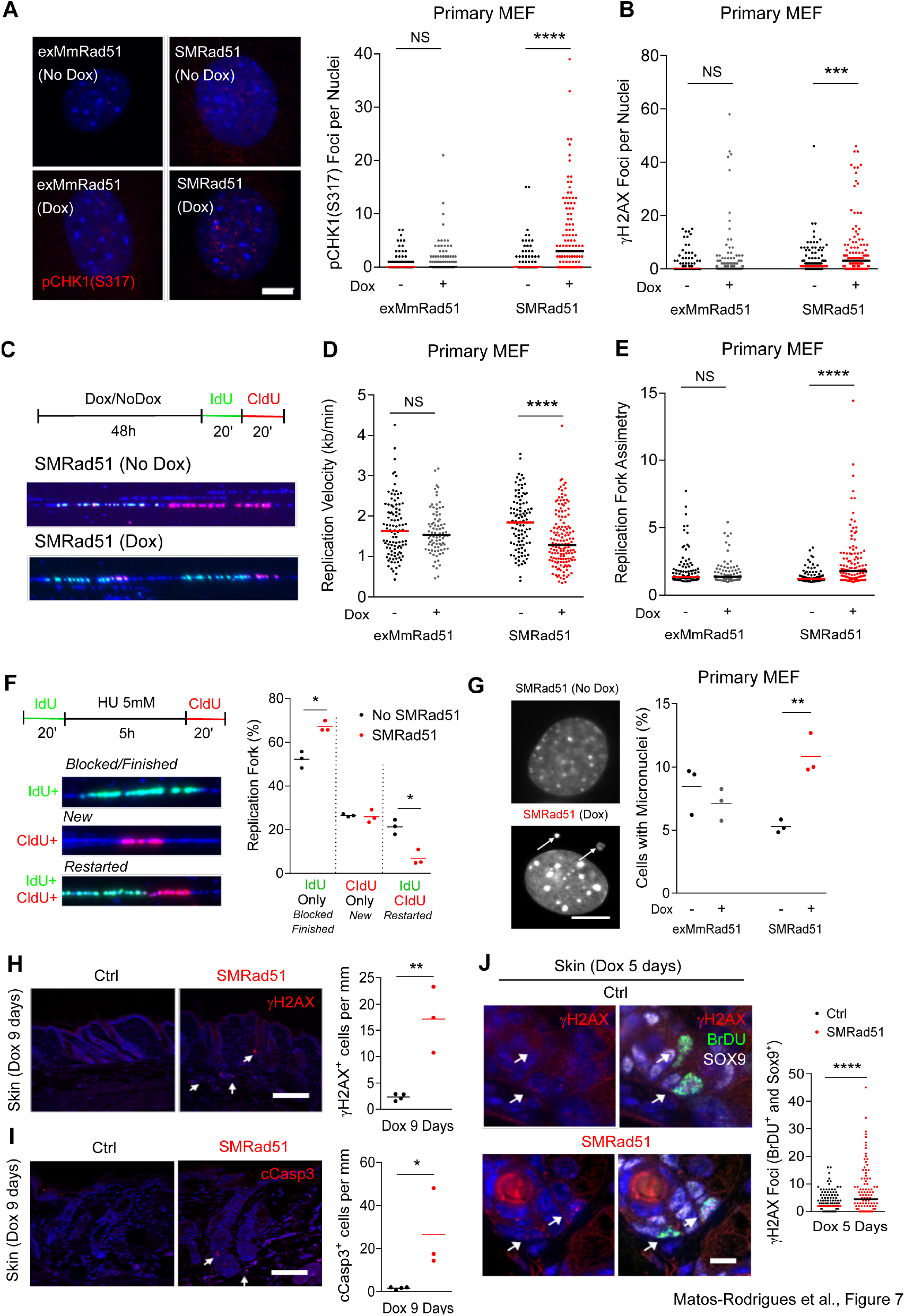
SMRAD51 leads to replication stress during unchallenged replication *in vivo*. **A.** Representative images (left panel) and quantification (right panel) of pChk1 (S317) foci in exMmRad51 and SMRad51 primary MEFs treated or not treated with Dox for 6 days (n=3 per group). **B.** Quantification of γH2AX foci per nucleus in exMmRad51 and SMRad51 primary MEFs treated or not treated with Dox for 6 days (n=4 per group). **C.** Representative scheme and pictures of molecular combing DNA fibers in SMRad51 primary MEFs treated or not treated with Dox (48 h). **D.** Quantitation of replication velocity analyzed by molecular combing in exMmRad51 and SMRad51 primary MEFs treated or not treated with Dox for 48 h. **E.** Replication fork asymmetry analysis by molecular combing in exMmRad51 and SMRad51 primary MEFs treated or not treated with Dox for 48 h. **F.** Analysis of fork restart. A representative scheme (upper panel) and a picture (bottom left panel) of replication restart after fork blockage by HU are shown. After one pulse of IdU (green), replication was blocked with 4 mM HU for 5 h. Fresh HU medium was added to remove the HU, and a pulse of CldU (red) was applied. Forks that did not restart did not incorporate CldU (green without red labeling), while forks that restarted incorporated both IdU and CldU (both green and red labeling). New replication initiation was not labeled by the initial IdU but incorporated CldU (red without green). Right panel: Quantitative analysis of the percentage of replication forks containing only IdU, only CldU or both markers (right) in immortalized MEFs expressing or not expressing *SMRad51*. The graphs represent three independent experiments. G. SMRAD51 expression induces micronucleus formation. Representative images (left) and quantification (right) of micronucleated cells after exMmRAD51 or SMRAD51 expression with Dox treatment (Dox +) or without Dox treatment (Dox -) for 6 days in primary MEFs are shown. White arrows: micronuclei. **H.** Analysis of γH2AX^+^ cells *in vivo*. Left panels: representative pictures. Right panel: quantification of γH2AX^+^ cells in skin sections of Ctrl and SMRad51 mice after 9 days of Dox treatment. **I.** Analysis of cCasp3 labeling *in vivo*. Left panels: representative pictures. Right panel: quantification of cCasp3^+^ cells in skin sections of Ctrl and SMRad51 mice after 9 days of Dox treatment. **J.** Representative images (left) and quantification (right) of γH2AX foci in BrdU^+^ (replicating) and Sox9^+^ (capillary bulb progenitor) cells (white arrow) in the skin sections of Ctrl and SMRad51 mice treated with Dox for 5 days. Statistical analysis: **(A, B, D, E, J)** Mann-Whitney test. ***p<0.001; ****p<0.0001; NS: nonsignificant. **(F, G, H, I)** Each point represents a biological replicate. Student’s t-test. * p<0.05; ** p<0.01. Scale bars: **(A)** 5 μm; **(H, I)** 100 μm; **(J)** 10 μm.

Replication stress often results in the accumulation of micronuclei. Consistent with the impact on replication stress shown above, *SMRad51* expression also led to an increase in the number of cells carrying micronuclei (Fig. 7*G*).

Taken together, the data demonstrate that SMRAD51 affects replication fork dynamics, resulting in replication stress in MEFs derived from our mouse model.

*In vivo*, as in adult mice (see Fig. 3), DNA damage (indicated by γH2AX staining) and apoptosis (indicated by cCasp3 staining) accumulated in young growing mice in different tissues upon *SMRad51* expression (Fig. 7*H*, 7*I* and *SI Appendix*, Fig.S7*A*, S7*B*). To evaluate whether SMRAD51 activates the DDR specifically in replicating cells *in vivo*, we quantified γH2AX foci in BrdU^+^ capillary bulb (Sox9^+^) progenitor cells in the skin of growing mice after 5 days of Dox treatment. *SMRad51* expression increased the numbers of γH2AX foci in BrdU^+^/Sox9^+^ (proliferating) cells (Fig. 7J), revealing that it induced replicative stress *in vivo*.

Altogether, these data show that RAD51 is essential for progenitor homeostasis *in vivo* and that its functional inactivation leads to replication stress and apoptosis, resulting in progenitor exhaustion in proliferative tissues and accounting for premature aging due to defective tissue renewal.

## Discussion

Here, the expression of the RAD51 dominant negative form *SMRad51*, which poisons the strand exchange activity of the RAD51-ssDNA filament (without activating nonconservative repair pathways), was found to induce premature aging and reduce lifespan. Remarkably, the expression of *SMRad51*, did not increase tumor prevalence in mice, although it generates genetic instability.

To circumvent limitations due to the fact that most HR genes are essential, we took advantage of a dominant negative form of RAD51, SMRAD51, that we previously designed (21, 28–32). HR competes with the non-conservative repair processes SSA and A-EJ (44–46). The binding of RAD51 to ssDNA not only triggers HR, but also prevents SSA and A-EJ, through DNA occupancy (21), thus preserving from the genetic instability that results from these nonconservative repair processes. Therefore, the knock out of RAD51 or the inhibition of HR mediator/accessory proteins such as BRCA2 or PALB2, which load RAD51 on ssDNA and/or stabilize the RAD51-ssDNA filament, results in the absence of RAD51 on ssDNA making it accessible to SSA and A-EJ pathways (21–23), which increase genomic instability. Therefore, the phenotypes of such mutants result from a mix of HR deficiency and increase of SSA and A-EJ. Importantly, because SMRAD51 binds damaged DNA (see *SI Appendix*, Fig. S2 and (21), its expression suppresses HR but without stimulation of the nonconservative repair mechanisms SSA and A-EJ (21). Thus, SMRAD51 is a unique tool enabling us to focus on the impact of the inhibition of solely HR, *in vivo*. Indeed, other dominant negative forms of RAD51, either mutated in Fanconi anemia group R (RAD51-T131P) or mutated in the ATP binding site (RAD51-K133R or RAD51-K133A) do not bind damaged DNA and impair the binding of endogenous WT-RAD51 *in vivo* (21, 47, 48); therefore, their expression leads to SSA and A-EJ stimulation (21). Here, we also further characterized the molecular mechanisms of HR inhibition by SMRAD51 (*SI Appendix*, Figs. S1 and S2). RAD51 promotes the central step of HR through homology searching and strand exchange with an intact DNA partner. This activity requires a well-ordered RAD51-ssDNA filament, which is thus the active species of HR. We show that by altering the structure of the RAD51-ssDNA filament, SMRAD51 inhibits RAD51 strand invasion activity, ultimately inhibiting HR in a dominant negative way. In our mouse model, we showed that SMRAD51 suppresses HR and generates replication stress, genomic instability, and systemic inflammation, ultimately resulting in premature aging. Consistent with the induction of replication stress, proliferating tissues and progenitors (which are replicating) are particularly affected. This should lead to progenitor pool exhaustion that hampers tissue renewal, thus leading to accelerated aging. Additionally, other systemic effects, such as inflammation, can aggravate the aging phenotype (Fig. 8). Collectively, the associations of defects in different tissues with inflammation can account for the reduced lifespan upon disruption of RAD51 function *in vivo*. Of note, these defects more strikingly affect young mice than adult mice, as most cells are dividing in young mice, thus replicating their genome,.

**Fig. 8.**
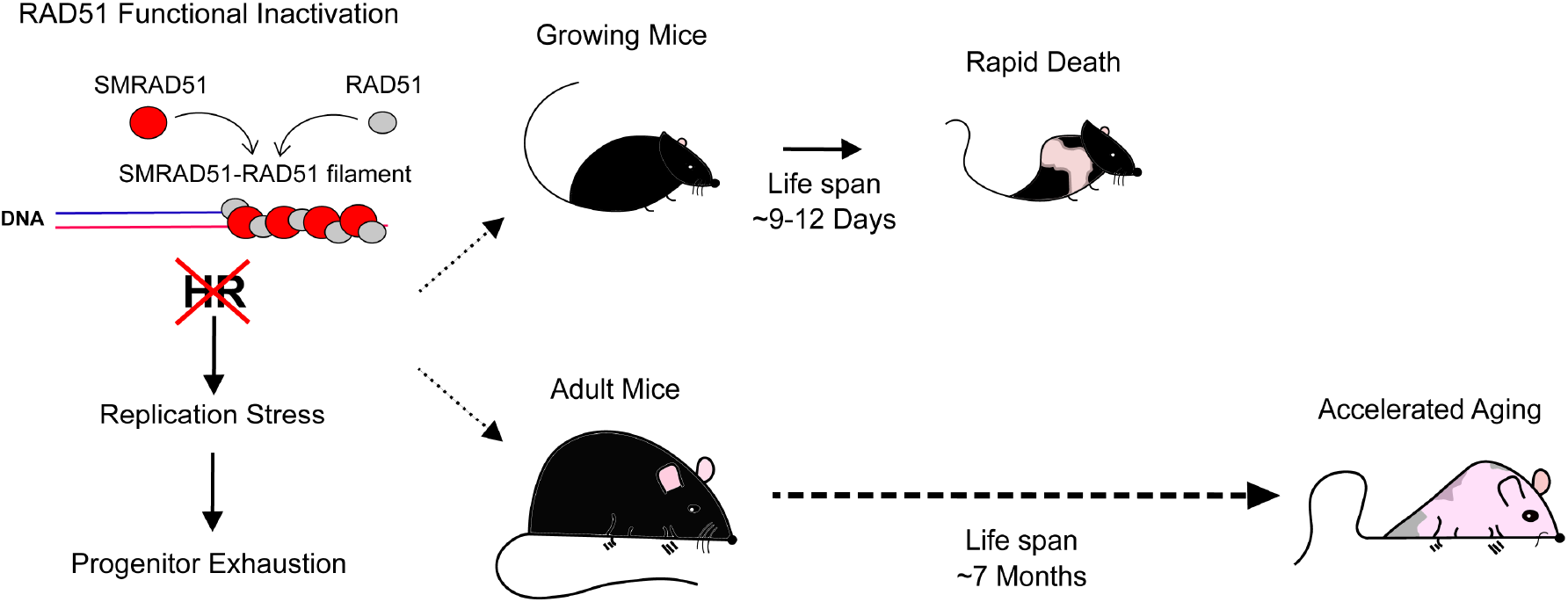
Impact of RAD51 functional inhibition *in vivo*. Expression of *SMRad51*, which alters the structure of the RAD51/ssDNA filament and inhibits HR (left panel), strongly affects mouse growth, hair and viability in growing young mice, leading to rapid mouse death. In adults, expression of *SMRad51* induces premature aging phenotypes and decreases lifespan.

According to the intrauterine programming model, developmental issues generated by replication stress during embryogenesis are the underlying cause of tissue degeneration and malfunction that results in a premature aging-like phenotype in adulthood (49, 50). Here, functional inactivation of RAD51 was performed in adults but still resulted in premature aging phenotypes. Therefore, the intrauterine programming model cannot account for the phenotypes observed in the current study. Instead, our results are consistent with data showing that genetic inactivation of the replicative stress response factor *Atr* leads to premature aging in adult mice (51).

Remarkably, in contrast with RAD51, mutations affecting factors that promote the loading of RAD51 onto ssDNA, such as BRCA1, BRCA2 and PALB2, confer cancer predisposition. Intriguingly, although RAD51 plays the pivotal role in HR, inactivating mutations of RAD51 have not been associated to carcinogenesis, although extensive studies have been performed, revealing the “RAD51 paradox” (13). Our data bring an experimental support to this paradox as, although SMRAD51 expression generated genetic instability, it did not favor tumorigenesis. One hypothesis is that mice with premature aging cannot support tumor development and die before tumors develop. A second hypothesis is that functional inactivation of RAD51 directly prevents the expansion of transformed cells and consequently tumor formation. Indeed, transformed cells are highly proliferative, and RAD51 functional inactivation preferentially affects highly proliferative cells. Third, tumorigenesis might be favored by the inhibition of HR associated with stimulation of mutagenic alternative pathways such as SSA and A-EJ, which rescue part of viability, but increase genome instability; RAD51 that protects against genetic instability, would protect both against tumor formation and premature aging but suppression of HR alone, without stimulation of SSA and A-EJ, leads to accelerated aging before tumor development (Fig. 8). Our data also provide evidence that, when premature aging is induced, genetic instability can be disconnected from oncogenesis.

## Materials and Methods

### Mice

*ExMmRad51* (HA-tagged *Mus musculus Rad51*)- and *SMRad51*-containing mice were generated using the following strategy. ExMmRad51 and SMRad51 were derived from previously developed plasmid constructs (30). Both transgenes were subcloned from a pcDNA3.1-puro plasmid containing exMmRad51 and SMRad51 to a pBI-4 plasmid containing the *tet* operators. The vectors were then linearized and electroporated into ES cells (129/SVEV). The clones were selected using puromycin and injected into blastocysts to enable germline transmission. For generation of the rtTA mouse line, mice containing the lox-stop-lox *rtTA-EGFP* transgene (B6.Cg-Gt(ROSA)26Sor^TM1(rtTA,EGFP)Nagy^/J) were mated with Vasa-Cre mice (FVB-Tg(Ddx4-cre)1Dcas/J) to obtain mice that constitutively expressed rtTA (these mice were therefore called rtTA mice). Littermates obtained from crosses of SMRad51 (*rtTA;SMRad51)* mice with control *(SMRad51)* mice or exMmRad51 *(rtTA;ExMmRad51)* mice with control (*exMmRad51)* mice were used for experimental analysis.

All mice were kept in a mixed genetic background. In young growing mice, *in vivo* transgene expression was induced by intraperitoneal injections of Dox diluted in PBS (5 μg/g, Sigma #D9891) on Monday, Wednesday and/or Friday. In adult mice, *in vivo* transgene expression was induced by *ad libitum* feeding of a diet with 625 mg/kg Dox (Sigma #D9891). All mice used as controls were treated with Dox.

The experimental procedures with animals were performed in accordance with French government regulations (Services Vétérinaires de la Santé et de la Production Animale, Ministère de l’Agriculture).

### Histology

Whole tissues or biopsies were fixed in 4% PFA immediately after dissection. The fixed tissues were then dehydrated and embedded in paraffin using a Tissue-Tek tissue machine (Sakura). The paraffin blocks were sectioned (5 μm) and placed on slides. Before staining, the sections were dewaxed and rehydrated. Pathological analyses were performed on hematoxylin (Path #10047105)-, eosin (Path #10047001)- and alcoholic saffron (Path #10047028)-stained sections of at least three different mice. Colorimetric images were captured using an Olympus BX51 microscope.

### Immunohistochemistry and immunofluorescence

For tissue immunostaining, the slides were washed with PBS, and antigen retrieval was performed using citrate buffer (pH=6). Antibodies were diluted in blocking buffer (5% NDS, 1% BSA and 0.1% Triton). For cultured cell immunostaining, cells were grown on glass coverslips and fixed with 4% PFA for 10 minutes at room temperature. The cells were then washed with PBS, permeabilized with 0.2% Triton and blocked with 1% BSA for 30 minutes. Primary antibodies were diluted in 1% BSA for immunostaining of cultured cells. The samples were incubated with the following primary antibodies overnight at 4°C: anti-SMRAD51 (directed against yeast N-terminal rad51) (1:100, Santa Cruz Biotechnology #SC33626), anti-Ki67 (1:100, BD Biosciences #556003), anti-SOX9 (1:100, Abcam #AB5535), anti-γH2AX (1:1,000, Bethyl #A300-081), anti-γH2AX (1:100, Millipore #05-636), anti-pCHK1 (S317) (1:100, Cell Signaling Technology #2344), anti-BrdU (1:100, Bio-Rad #OBT0030G), anti-PLZF (1:50, Santa Cruz #SC22839), anti-cCasp3 (1:100, Cell Signaling Technology #9661S) and anti-HA (1:200, Santa Cruz #7392). For immunofluorescence, the samples were incubated with Alexa-conjugated secondary antibodies for 2 h at room temperature, and the slides were mounted with DAPI ProLong Gold (Thermo Fisher #P36931). For immunohistochemistry, the slides were incubated for 30 minutes with HRP-conjugated secondary antibodies, and the staining was then revealed with DAB (Vector #SK4100). Fluorescence images were captured using a Nikon Eclipse Ti2 microscope, and colorimetric images were captured using an Olympus BX51 microscope.

To label S-phase cells *in vivo*, we performed intraperitoneal BrdU injection (50 μg/g of body weight) (Sigma Aldrich #B5002). Samples were collected 1 h after injection for analysis.

### Blood and bone marrow analysis

Bone marrow cells from the femur and tibia were flushed with PBS. After hemolysis of blood or bone marrow samples, the cells were counted using an Abbott Cell Dyn 3700 machine. The cells were then labeled using the following antibodies: anti-Lin^+^ (MACS), anti-c-Kit (BioLegend), anti-Sca1 (Thermo Fisher), anti-IL7R (BD Bioscience), anti-CD3 (BioLegend), anti-B220 (Thermo Fisher), anti-Gr1 (BioLegend), and anti-FLK2 (Thermo Fisher). Data acquisition was performed on a FACSCanto II (BD Biosciences), and the data were analyzed with FlowJo software.

### MEFs

MEFs were isolated from embryonic day 12.5 (E12.5) embryos and immortalized with SV40 (Addgene plasmid #21826). Primary MEFs were used up to passage 4. Immortalized MEFs (iMEFs) and primary MEFs were cultured in DMEM (Gibco #41965-039) supplemented with 10% tetracycline-free FBS (Takara #631107) and penicillin-streptomycin (Gibco #15140122).

### RNA extraction, cDNA synthesis and real-time RT-PCR

RNA was extracted from skin biopsies (~1 cm^2^) taken from the upper backs of mice (Qiagen #74106). Each RNA sample was collected in 40 μL of ultrapure water (Thermo Fisher #10977). The RNA concentration and purity were determined using a NanoDrop spectrophotometer (Eppendorf). The RNA integrity was analyzed by 1% agarose gel electrophoresis. cDNA was generated using a high-capacity cDNA synthesis kit (Thermo Fisher #4368814) following the manufacturer’s instructions.

We used 384-well optical plates to perform real-time RT-PCR using an Applied Biosystems 7900HT Fast Real-Time PCR System thermocycler. The primer sequences used for real-time RT-PCR are shown in *SI Appendix*, Table S2. The real-time RT-PCR mix included 6.25 μL of 2× SYBR Green mix, 2 μL of diluted cDNA (1:10), 0.25 μL (5 μM) of each primer and 4.25 μL of ultrapure water (Gibco #10977). The cycling conditions included 50°C for 2 minutes, 95°C for 10 minutes and 40 cycles of 95°C for 15 s and 60°C for 60 s. The delta-delta Ct (2-ΔΔCt) method was used to determine the relative quantities of the target genes compared to the reference genes. We used the average of the two reference genes (*Gapdh* and *β-actin*) as the final reference.

### Cytokine array

For protein array analysis, approximately 150 μL of blood serum containing protease inhibitors (Thermo Fisher #78438) was used. The serum was stored at −80°C until analysis. Samples from two mice of each genotype (Ctrl or SMRad51) that had been fed a Dox-containing diet for 3 months were used. A Proteome Profiler Mouse XL Cytokine Array Kit (R&D Systems #ARY028) was used following the manufacturer’s instructions. In *SI Appendix*, Table S3 (adult mice) and *SI Appendix*, Table S4 (young growing mice), we present the results of densitometry analysis and the comparisons.

### Replication dynamics analysis

For a DNA spreading assay, replicating DNA in immortalized MEFs was labeled using 50 μM CldU and 50 μM IdU for 20 minutes each. Cells were then harvested and resuspended in cold PBS. DNA spreads were performed using 1×10^3^ cells. To extend DNA fibers, 2 μL of cells were incubated for 3 minutes with 7 μL of lysis buffer (0.5% SDS, 50 mM EDTA, 200 mM Tris-HCl) in the upper parts of microscope slides. To generate single-DNA-molecule spreads, the slides were turned at a 15° angle, which allowed the genomic DNA to spread by gravity. The DNA fibers were then fixed in methanol and acetic acid (3:1) and stored at 4°C.

For molecular combing, a single assay was performed as previously described ((52)). Briefly, primary MEFs were incubated or not incubated with Dox for 48 h and pulse-labeled with IdU for 20 minutes and then with CldU for 20 minutes. The cells were collected, and the DNA fibers were purified by enzymatic protein digestion in agarose blocks and subsequently stretched at a rate of 2 kb/μm on silanized coverslips (52).

Immunofluorescence detection for the DNA combing and DNA spreading experiments was performed with the following antibodies, in order: (1) mouse anti-BrdU (BD Biosciences #347583) and rat anti-BrdU (AbD Serotec #OBT 0030), (2) A488-conjugated goat anti-mouse (Invitrogen #A11029) and A555-conjugated goat anti-rat (Abcam #A21434), (3) mouse anti-ssDNA (Millipore #MAB3034), (4) Cy5.5-conjugated goat anti-mouse (Abcam #ab6947) and (5) Cy5.5-conjugated donkey anti-goat (Abcam #ab6951). Fluorescence images were acquired using an epifluorescence microscope (AxioImager Z2; Carl Zeiss). MetaMorph software (Roper Scientific) was used to acquire images and to build a large-scale mosaic of up to 100 images. This technique theoretically enabled us to recover long fibers up to approximately 3 Mb in length. We systematically used ssDNA staining to ensure that the replication signals belonged to the same fiber.

### Western blot analysis

Proteins were extracted using RIPA buffer (Thermo Fisher #89900) supplemented with phosphatase inhibitors (Thermo Fisher #P0044 and #P5726) and a protease inhibitor (Thermo Fisher #78438). After SDS-PAGE protein separation, the proteins were transferred to a nitrocellulose membranes overnight at 30 V and 4°C. The membranes were incubated with the following primary antibodies, which were diluted in 5% milk prior to incubation: anti-RAD51 (1:1,000, Millipore #PC130), anti-STAT1 (1:10,000, Cell Signaling Technology #9164), anti-pSTAT1 (Tyr701) (1:10,000, Cell Signaling Technology #14994), anti-VINCULIN (1:10,000, Abcam #AB18058) and anti-HA (1:1,000, BioLegend #MMS-101R). HRP-conjugated secondary antibodies were purchased from Thermo Fisher (1:10000, anti-mouse IgG, #31430, anti-rabbit IgG, #31460). A Pierce (Thermo Fisher #32106) or Luminata (Thermo Fisher #WBLUF0500) ECL system was used according to the manufacturer’s instructions, and chemiluminescence was captured using an Amersham Imager 600 (GE Life Sciences).

### Statistical analysis

GraphPad Prism software was used for statistical analysis. One- or two-way ANOVA was performed as indicated. The p-values are based on two-sided tests.

## Acknowledgments

We thank George Garinis for helpful and stimulating discussion and Didier Busso and Xavier Veaute from the CIGEX (CEA, Fontenay aux Roses) for providing us the purified SMRAD51 and MmRAD51 proteins.

