## supplementary material for "*In vivo* inactivation of RAD51-mediated homologous recombination leads to premature aging, but not to tumorigenesis"

**This PDF file includes:**

Supplementary text
Figures S1 to S7
Tables S1 to S2
Additional material and methods
SI References

**Other supplementary materials for this manuscript include the following:**

Table S3 to S5 (see excel tables)
Movies S1 to S2

### Supplementary Information Text

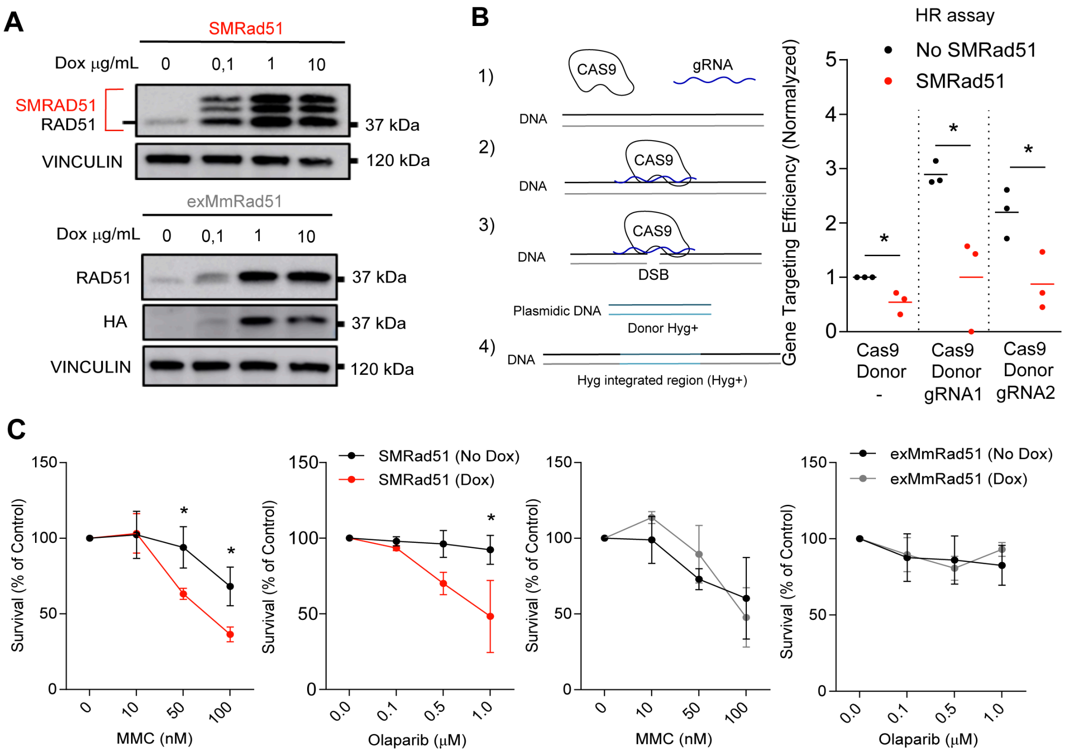


**D**

Genetic modifications in cells treated with Dox compared to cells without treatment

| Genotype | SNV (blue dots) | Deletions (red lines) |
|---|---|---|
| Ctrl | 16,8 per genome | 6,8 per genome |
| SMRad51 | 45,15 per genome | 9,52 per genome |

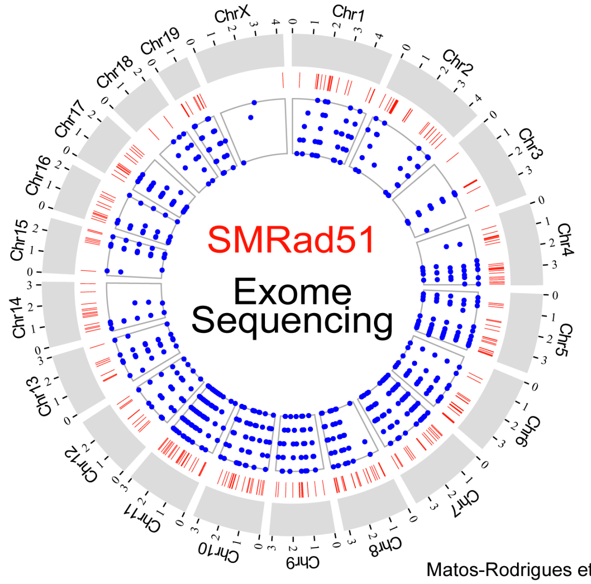


Matos-Rodrigues et al., Supplementary Figure 1

**Fig. S1. *SMRad51* expression decreases HR efficiency, increases sensitivity to genotoxic agents in immortalized MEFs and leads to genomic instability in primary MEFs.** (**A**) Transgene expression as assessed by Western blot analysis in immortalized exMmRad51 or

SMRad51 MEFs treated or not treated with increasing concentrations of Dox for 24 h. SMRAD51 expression resulted in 3 characteristic diagnostic bands (1–5). (**B**) Homology-directed repair-mediated gene targeting assay. The protocol consisted of sequence homology-dependent replacement of the second exon of the 53BP1 gene by a hygromycin resistance (Hyg+) gene using homologous donor DNA (here called Donor) after Cas9-mediated induction of a DSB in the targeted sequence (left panel). Cells with and without SMRad51 expression were transfected with three combinations of plasmids: 1) Cas9+Donor (no Cas9 activity) and 2) Cas9+Donor+gRNA1 (Cas9 active) or 3) Cas9+Donor+gRNA2 (Cas9 active). Recombinant cells became resistant to hygromycin. Right panel: Quantification from 3 independent experiments. (**C**) Quantification of cell viability by colony formation assay for immortalized MEFs expressing or not expressing SMRad51 or exMmRad51 observed 10 days after MMC or Parp inhibitor (olaparib) treatment. Dox treatment was started 24 h before exposure to genotoxic drugs. (**D**) Comparison of genomic instability by whole-exome sequencing after Dox treatment in SMRad51 or Ctrl cells treated with Dox (10 μg/mL) for 6 days. The Circos plot shows the positions of mutations (blue dots) and deletions (red lines) and in the chromosomes (Chrs) of SMRad51 primary MEFs. The analyses are presented in Table S5. Statistical analysis: (b) Student's t-test. (**C**) Two-way ANOVA followed by Sidak posttest, Student's t-test. *$p<0.05$; **$p<0.01$; ***$p<0.001$. The error bars represent the ±SEM.

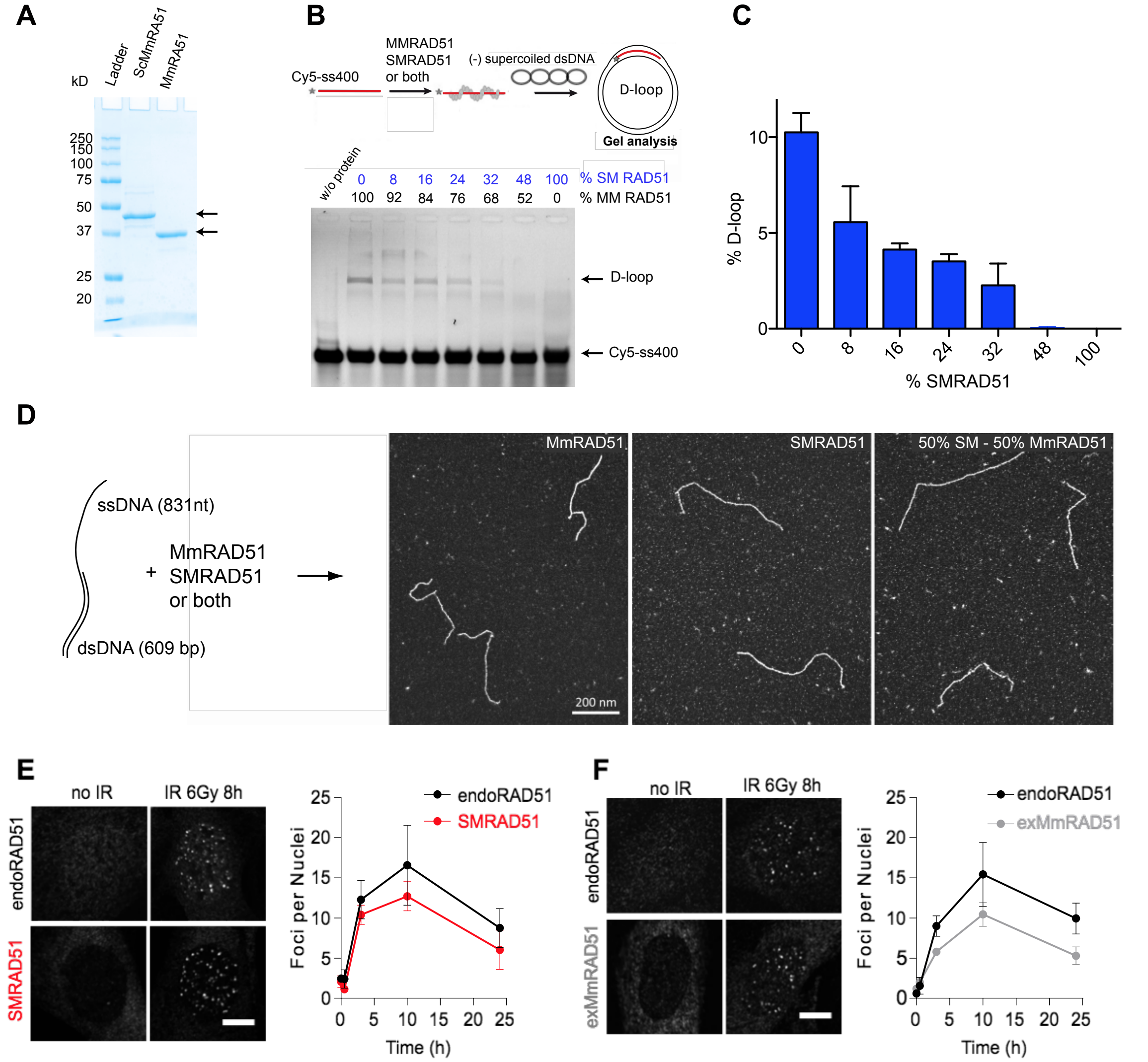


**Fig. S2. Cellular and biochemical characterization of SMRAD51. (A)** Electrophorese analysis of the purified SmRAD51 and MmRAD51 proteins. (**B**) Impact of SMRAD51 on the MmRAD51-mediated D-loop assay. Upper panel: scheme of the D-loop assay. Lower panel: electrophoresis gel of the D-loop formation experiment. Lane 1: in the absence of proteins; other lanes: different mix of MmRAD51 and SMRAD51 proteins. The respective percentages of each protein are indicated. (**C**) Quantification of the D-loop assay. (**D**) Transmission electron microscopy (TEM) analysis. Left panel: the DNA substrate. Right panels: representative pictures of presynaptic filaments formed by MmRAD51, SMRAD51 or both. Note that &00% of DNA molecules were coated by MmRAD51 or SMRAD51. Scale bar: 100 nm. (**E, F**) Representative pictures and quantification of endogenous MmRAD51 (endoRad51), SMRAD51 and exMmRAD51 focus formation dynamics after irradiation. Immortalized MEFs were treated or not treated with Dox for 24 h, exposed to ionizing irradiation (6 Gy) and fixed at different time points. SMRAD51 was stained with a specific antibody. exMmRAD51 was stained with an HA-tag antibody. The graph shows the ±SEMs of three independent experiments. EndoRAD51 staining was performed in cells not treated with Dox and stained for RAD51. Scale bars: 5 μm.

**A**

| Organ | Phenotype (SMRad51) |
|---|---|
| Kidney | Normal histology |
| Stomach | Normal histology |
| Pancreas | Normal histology |
| Heart | Normal histology |
| Liver | Normal histology |
| Large Intestine | Normal histology |
| Small Intestine | Normal histology |
| Ovary | Normal histology |
| Spleen | Presence of hematopoietic figures |
| Testicules | Reduced size with normal histology |
| Lung | Edematous alveolitis |
| Skin | Squamous epithelium with hyperplasia, atypia and hyperkeratosis (ortho). 1 of 6 mice presented squamous cell carcinoma. |

**B**

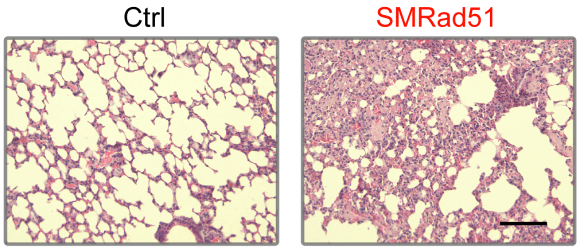


**Fig. S3. Histopathological analysis of adult mice after *SMRad51* expression for 3 months. (A)** Table showing a summary of the pathological analysis results for various tissues of Ctrl and SMRad51 adult mice fed a Dox-containing diet for 6 months. At least 3 different mice of each genotype were analyzed. **(B)** Representative images of hematoxylin-eosin-saffron staining in lungs. Scale bar: 100 μm.

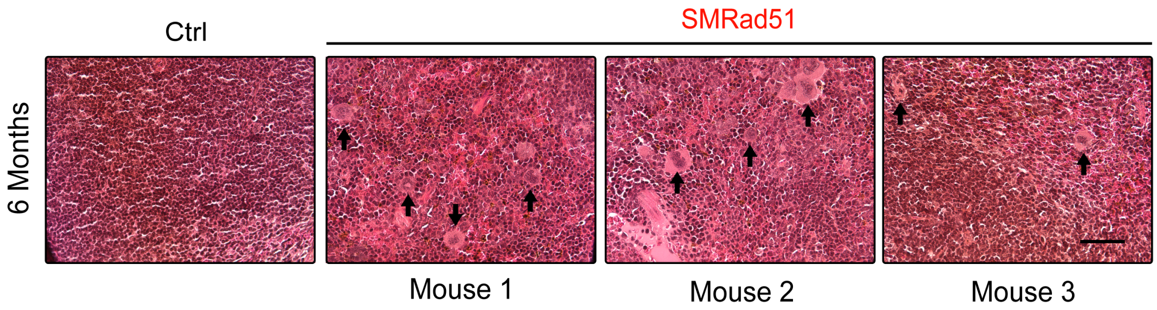


**Fig. S4. Expression of *SMRad51* in adult mice leads to extramedullary hematopoiesis in the spleen.** Representative images of hematoxylin-eosin-saffron staining in spleen sections of Ctrl and SMRad51 adult mice fed a Dox-containing diet for 6 months. Arrows: overrepresentation of megakaryocytes in the spleens of *SMRad51*-expressing mice.

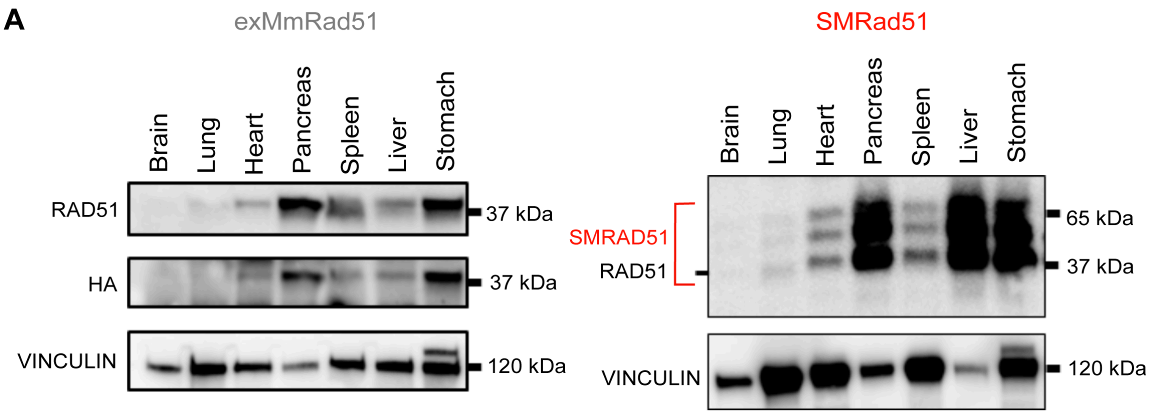


**B**

| Organ | Phenotype | SMRAD51 expression |
|---|---|---|
| Pancreas | Normal histology | Yes |
| Heart | Normal histology | Yes |
| Lung | Normal histology | Yes |
| Liver | Normal histology | Yes |
| Spleen | Normal histology | Yes |
| Small intestine | Normal histology | Yes |
| Testicules | Features of arrested development | Yes |
| Skin | Few and abnormal hair folicules, acanthosis, numerous inflammatory cells | Yes |

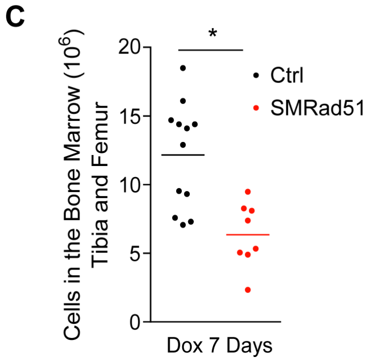


**Fig. S5. Histopathological analysis of young growing mice after expression of *SMRad51* for 12 days. (A)** Western blot analysis of exMmRAD51 and SMRAD51 protein content in extracts from various tissues of young mice after 5 days of Dox treatment. **(B)** Summary of the results of pathological analysis and SMRAD51 presence assessment by immunohistochemistry in various tissues of SMRad51 and Ctrl mice. At least 3 different mice of each genotype were analyzed. **(C)** Cell numbers in tibia and femur bone marrow samples from young Ctrl and SMRad51 mice after 7 days of Dox treatment. Each point represents a biological replicate. Statistical analysis: Student's t-test. *$p<0.05$.

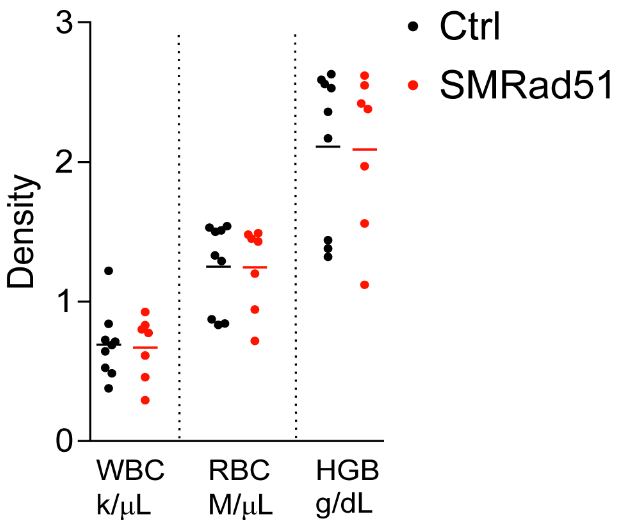


**Fig. S6. *SMRad51* expression for 7 days does not cange basic blood parameters**. WBC, RBC and HGB density analysis of blood samples from growing mice after 7 days of Dox treatment. Each point represents a biological replicate.

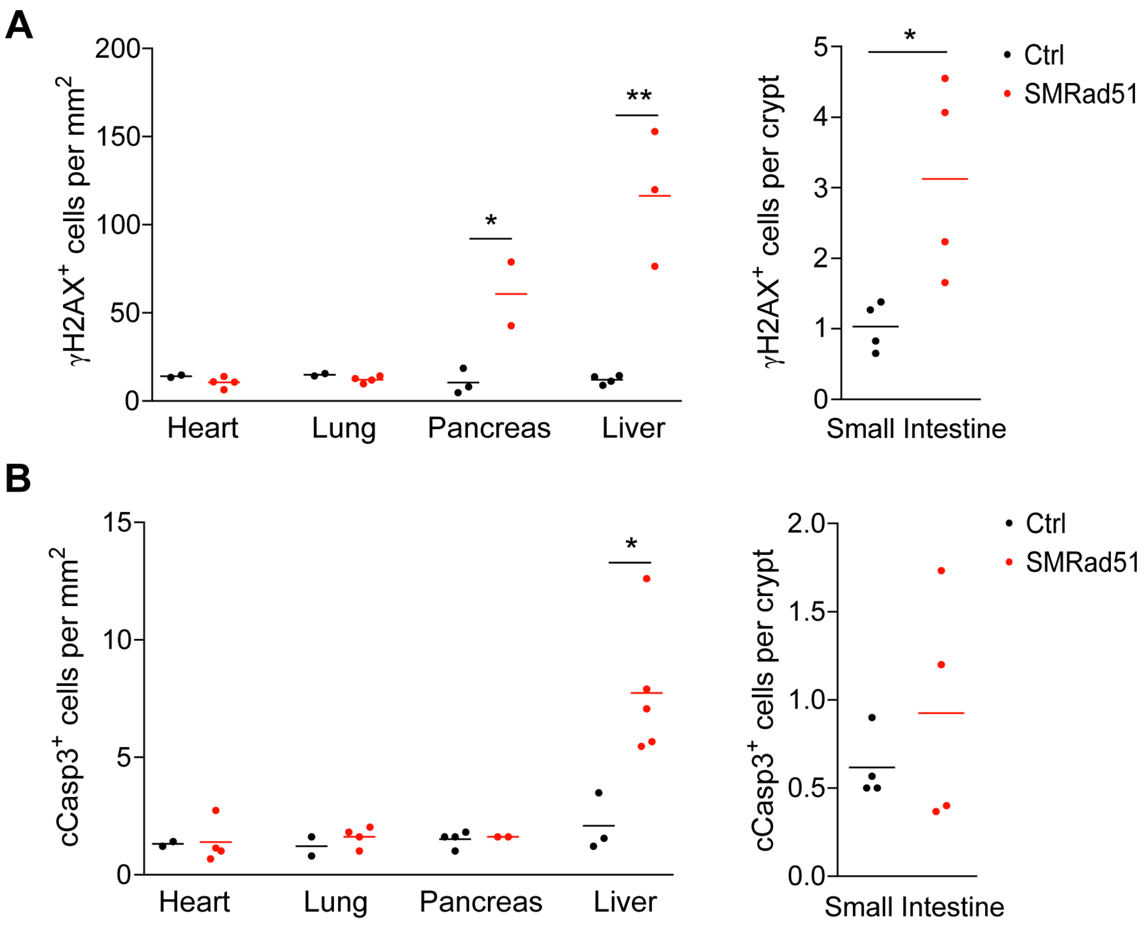


**Fig. S7. *SMRad51* expression in growing mice leads to tissue-specific DDR activation and apoptosis.** (A) Quantification of γH2AX⁺ cells in various tissues of Ctrl and SMRad51 mice after 12 days of Dox treatment. (B) Quantification of cCasp3⁺ positive cells in various tissues of Ctrl and SMRad51 mice after 12 days of Dox treatment. Each point represents a biological replicate. Statistical analysis: Student's t-test. * $p<0.05$; ** $p<0.01$.

**Table S1.** Pre-mature ageing phenotypes compared to other models of DNA repair and/or progeroid syndrome

| Phenotype | *ERCC1*$^{□/-}$ | *XPD*$^{TTD}$ | *mTR*$^{-/-}$ | *ATR*$^{s/s}$ | *BRCA1*$^{Δ11}$*53BP1*$^{S25A}$ | SMRad51 | ExMmRad51 | *Lmna*$^{HG/+}$ |
|---|---|---|---|---|---|---|---|---|
| | (6–8) | (9, 10) | (11) | (12) | (13) | | | (14) |
| **Hair Loss** | - | + | + | + | na | + | - | na |
| **Kyphosis** | + | + | + | + | + | + | - | + |
| **Reduced Activity** | + | na | na | na | + | + | - | na |
| **Priapism** | + | na | na | na | na | + | - | na |
| **Decreased Body weight** | + | + | + | + | + | + | - | + |
| **Subcutaneous Fat Loss** | + | + | + | na | na | + | - | + |

**Table S2.** List of primers used for realtime RT-PCR.

| Gene Symbol | Forward primer 5' - 3' | Reverse primer 5' - 3' |
|---|---|---|
| *β-Actin* | GCCCTGAGGCTCTTTTCCAG | TGCCACAGGATTCCATACCC |
| *Gapdh* | AGGTCGGTGTGAACGGATTG | TGTAGACCATGTAGTTGAGGTCA |
| *Il6* | TAGTCCTTCCTACCCCAATTTCC | TTGGTCCTTAGCCACTCCTTC |
| *Ccl2* | TTAAAAACCTGGATCGGAACCA<br>A | GCATTAGCTTCAGATTTACGGGT |
| *Il1β* | CAGGCAGGCAGTATCACTCA | AGCTCATATGGGTCCGACAG |
| *Tnfα* | GAACTGGCAGAAGAGGCACT | AGGGTCTGGGCCATAGAACT |
| *Cxcl1* | CTGGGATTCACCTCAAGAACAT<br>C | CAGGGTCAAGGCAAGCCTC |

### Additional material and methods

#### *Colony formation assay*

For a colony formation assay, one hundred exMmRad51 or SMRad51 iMEFs were plated per well in 6-well plates. After 24 h, the cells were treated or not treated with 10 µg/mL doxycycline, and 24 h later, the cells were treated or not with mitomycin C (MMC; Roche #10107409001) or olaparib (Selleckchem #S1060) for 10 days. The clones were stained with crystal violet and counted.

#### *Homology-dependent gene targeting assay*

The gene targeting procedure was derived from (15). This protocol consists of replacing the second exon of the 53BP1 gene with a hygromycin resistance ($Hyg^+$) gene using a homology-dependent donor plasmid (here called pDonor) after Cas9-mediated production of a DNA DSB. A total of 300,000 cells were resuspended in 100 µL of nucleofection buffer (Lonza #VCA-1003) containing 1 µg of pCas9 and 2 µg of pDonor with or without 1 µg of pgRNA1 or pgRNA2. The cells were nucleofected using Amaxa (Lonza) protocol T30. Immediately after nucleofection, 500 µL of medium was added, and the cells were plated on a 10 mm petri dish. After 48 h, the targeted clones ($Hyg^+$) were selected for 10 days with medium containing hygromycin that was renewed every 48 h. The clones were stained with crystal violet and counted.

#### *Protein purification*

Previously generated pcDNA3.1-MmRad51 and pcDNA3.1-SMRad51 constructs (16) encoding sequences were subcloned into a pCDF-His-SUMO backbone using a complementary single-strand annealing-based method. His-SUMO-MmRAD51 and his-SUMO-SMRAD51 were both expressed in the *E. coli* strain BRL(DE3)pLysS. All of the protein purification steps were carried out at 4°C. Protein expression was induced in 3 L of cell culture medium with 0.5 mM isopropyl-1-thio-β-D-galactopyranoside overnight at 20°C, after which the cells were resuspended in PBS with 350 mM NaCl, 20 mM imidazole, 10% glycerol, 0.5 mg/mL lysozyme, complete protease inhibitor (Roche), and 1 mM 4-(2-aminoethyl)benzenesulfonyl fluoride (AEBSF). Then, the cells were lysed by sonication, and the insoluble material was removed by centrifugation at 150,000 × *g* for 1 h. The supernatant was incubated with 5 mL of Ni-NTA resin (Qiagen) for 2 h. The mixture was poured into an Econo-Column Chromatography Column (Bio-Rad), and the beads were washed with 80 mL of W1 buffer (20 mM Tris HCl pH 8, 500 mM NaCl, 20 mM imidazole, 10% glycerol, 0.5% NP40) followed by 80 mL of W2 buffer (20 mM Tris HCl pH 8, 100 mM NaCl, 20 mM imidazole, 10% glycerol, 1 mM DTT). His-SUMO-tagged proteins bound to the beads were then resuspended in 8 mL of W2 buffer and incubated with SUMO protease at a ratio of 1/80 (w/w) for 16 h. The proteins in which the His-SUMO tag was cleaved were then recovered in the flowthrough and directly loaded onto a HiTrap heparin column (GE Healthcare). The column was washed with W2 buffer, and then a 0.1-1 M NaCl gradient was applied. Fractions containing purified proteins were dialyzed against storage buffer (20 mM Tris HCl pH 8, 50 mM KCl, 0.5 mM EDTA, 10% glycerol, 1 mM DTT, 0.5 mM AEBSF), aliquoted and stored at -80°C. The concentrations of purified MmRad51 and SMRad51 were calculated using extinction coefficients of $1.664 \times 10^4$ $M^{-1}cm^{-1}$ and $1.813 \times 10^4$ $M^{-1}cm^{-1}$ at 280 nm, respectively.

#### *D-Loop assay*

For the D-loop assay, 19 nM molecules (7.5 µM nt) of a 400 nulcetides ssDNA labeled with the Cy5 fluorophore were incubated with MmRAD51, SMRAD51 or both (SM:MmRAD51 ratio: 0, 8, 16, 24, 32, 48 and 100%) at a final concentration of 2,5 µM (1 protein per 3 nt) for 3 minutes at 37°C. RPA was then added to a final concentration of 0.075 µM (1 protein per 100 nt) over a span of 15 minutes in buffer containing 10 mM Tris-HCl (pH 8), 50 mM sodium chloride, 2 mM calcium chloride, 2 mM Magnesium chloride, 1 mM DTT and 1.5 mM ATP. Then, a homologous double-stranded DNA (dsDNA) pUC19 plasmid (purchased from New England Biolabs) was added to the reaction at a concentration of 16 nM molecules in a final volume of 8 µL over a span of 30 minutes. The total reaction was stopped by the addition of 1% SDS (w/v) plus 25 mM EDTA and deproteinized (by 30 minutes of incubation at 37°C with 2 mg/mL proteinase K). The samples were run in a 1.2 % (w/v) agarose gel at 80 V for 35 minutes in 0.5× TAE buffer. Fluorolabeled DNA species were visualized by using a Typhoon FLA 9500 (GE Healthcare Life Sciences).

***TEM analysis of RAD51 filaments***

RAD51 presynaptic filaments were assembled on a 5' DNA junction with a single-stranded overhang (the construction of which has been described in ((17))) using MmRAD51, SMRAD51 or a mix of both, as follows: 0.2 µM (36 µM nt) of 5' junction DNA molecules were incubated with MmRAD51, SMRAD51 or both at a final concentration of 12 µM (1 protein per 3 nt) for 5 minutes at 37°C. RPA was then added to a final concentration of 0.6 µM (1 protein per 100 nt) over a span of 10 minutes in buffer containing 10 mM Tris-HCl (pH 8), 50 mM sodium chloride, 2 mM calcium chloride, 1 mM DTT and 1.5 mM ATP. The reactions were quickly diluted 25× in buffer, and a 5 µL drop was deposited on a 600-mesh copper grid previously covered with a thin carbon film and activated with pentylamine by glow discharge using a Dubochet device. Each grid was rinsed, positively stained with aqueous 2% (w/v) uranyl acetate, dried carefully with filter paper and observed in annular dark-field mode using a Zeiss 902 transmission electron microscope. Images were captured at a magnification of 85,000× with a MegaViewIII charge-coupled device (CCD) camera and analyzed with iTEM software (Olympus Soft Imaging Solution).

***Whole-exome sequencing***

Whole-exome sequencing was performed and analyzed with the genomic and bioinformatics platforms of the Institute Gustave Roussy. SMRad51 and control primary MEFs were treated or not treated with 10 µg/mL Dox for 6 days. DNA extraction and purification were performed via a standard phenol-chloroform method and RNase treatment. DNA quality was estimated with a Bioanalyzer (Agilent). Two hundred nanograms of genomic DNA was sheared with the Covaris S220 system to obtain a 150-base Gaussian distribution using a peak power of 140, a duty factor of 10, and a cycle/burst setting of 200 (LGC Genomics/KBioscience). DNA fragments were end-repaired, extended with an 'A' base on the 3′ end, ligated with paired-end adaptors with the Bravo Platform (Agilent) and amplified. Adaptor-ligated libraries were hybridized for 40 h with biotinylated oligo RNA baits using murine SureSelect capture probes (Agilent) and enriched with streptavidin-conjugated magnetic beads. The final libraries were indexed, quantified by real-time RT-PCR, pooled and sequenced using the default cluster method in paired-end sequencing mode (2×100 bp reads) on an Illumina NovaSeq 6000 sequencer. The mean depth in the targeted regions was 110× for all samples. The reads were mapped to the reference genome mm10 using the Burrows–Wheeler Aligner (BWA) alignment tool version 0.7.10. PCR duplicates were removed using SAMtools version 1.9. Local realignment around indels and base quality score recalibration were performed using the Genome Analysis Tool Kit (GATK) 3.7. Somatic single-nucleotide variations (SNVs) and indels were called using VarScan (v2.3.9). Variants were adopted as candidate mutations when the somatic P value was <0.001 and when the allele frequency was <0.01 in the reference sample and >0.01 in the treated sample. Additionally, variants with a read depth less than 50 in both reference and treated samples were filtered out. The variants were annotated with ANNOVAR version 20180416. We excluded SV129 strain-specific polymorphisms using the results from the Mouse Genomes Project release v5: REL-1505-SNPs_Indels. Structural rearrangements were identified using Manta version 1.5.0 in exome mode with the default parameters. Variations greater than 500 bp were considered structural rearrangements. A summary of the data is presented in *SI Appendix*, Table S5. A graphical representation of exome sequencing was created using the BioCircos R package. The whole-exome sequence ng ArrayExpress accession number is E-MTAB-8625.

***Data and materials availability:*** Whole exome sequencing ArrayExpress accession number: E-MTAB-8625.

**Movie S1 (separate file).** SupplmantaryVideo S1 SMRAD51.mov.

**Movie S1 (separate file).** SupplmantaryVideo S2_Control.mov

**Dataset Table S3 (separate file).** Table S3-Cytokine arrays in adult mice .xlsx

**Dataset Table S4 (separate file).** Table S4-Cytokine arrays in young mice.xlsx
**Dataset Table S5 (separate file).** Table S5-genetic alteration upon SMRAD51 expression.xlsx
